# Cell-cell metabolite exchange creates a pro-survival metabolic environment that extends lifespan

**DOI:** 10.1101/2022.03.07.483228

**Authors:** Clara Correia-Melo, Stephan Kamrad, Christoph B. Messner, Roland Tengölics, Lucía Herrera-Dominguez, St John Townsend, Mohammad Tauqeer Alam, Anja Freiwald, Kate Campbell, Simran Aulakh, Lukasz Szyrwiel, Jason S. L. Yu, Aleksej Zelezniak, Vadim Demichev, Michael Muelleder, Balázs Papp, Markus Ralser

## Abstract

Metabolism is fundamentally intertwined with the ageing process. We here report that a key determinant of cellular lifespan is not only nutrient supply and intracellular metabolism, but also metabolite exchange interactions that occur between cells. Studying chronological ageing in yeast, we observed that metabolites exported by young, exponentially growing, cells are re-imported during the stationary phase when cells age chronologically, indicating the existence of cross-generational metabolic interactions. We then used self-establishing metabolically cooperating communities (SeMeCos) to boost cell-cell metabolic interactions and observed a significant lifespan extension. A search for the underlying mechanisms, coupling SeMeCos, metabolic profiling, proteomics and genome-scale metabolic modelling, attributed a specific role to methionine consumer cells. These cells were enriched over time, adopted glycolytic metabolism and increased export of protective metabolites. Glycerol, in particular, accumulated in the communal metabolic environment and extended the lifespan of all cells in the community in a paracrine fashion. Our results hence establish metabolite exchange interactions as a determinant of the ageing process and show that metabolically cooperating cells shape their metabolic environment to achieve lifespan extension.

## Introduction

Metabolism is interlinked with the ageing process and determines lifespan at multiple levels. Processes driven or dependent on metabolism include cellular growth, ageing and death, energy production and the formation of molecular building blocks required for cellular homeostasis and repair, and even regulation of antimicrobial responses ^1–6^. Moreover, metabolic sensing and metabolic signalling systems that regulate cell growth and energy expenditure, such as the AMPK, Sirtuin and mTOR pathways, are also central pathways regulating cellular ageing and lifespan ^7–11^. Indeed, metabolism is a source of both damaging and protective molecules, and hence strongly intertwined with ageing processes. For example, metabolites like NADPH and glutathione fuel the metabolic antioxidant machinery and protect cells from oxidative damage that occurs during cellular ageing ^12^. On the other end of the spectrum, reactive products of metabolism such as methylglyoxal or superoxide, indiscriminately react and damage cellular membranes, proteins and nucleic acids, accelerating cellular ageing phenotypes ^13–15^. As a consequence, metabolic activity that determines the equilibrium between the levels of protective and damaging reactive molecules inside a cell is a critical determinant of ageing and lifespan ^16, 17^.

Cellular metabolism not only occurs inside cells, it also involves the exchange of metabolites between cells and tissues ^18–22^. In microbes, metabolite exchange interactions are a result of cells exporting metabolites into their surrounding and re-importing metabolites produced by themselves or neighbouring cells. Metabolite exchange interactions can be costly for cells, because the exported metabolites can be lost to competitors or diffusion ^23^, but are critical for cell growth and emerge for at least three reasons. First, the ability to uptake and exploit metabolites that are available environmentally renders cells more competitive. This means cells profit from possessing metabolite uptake systems even in situations where there are no advantages to be gained from cell-cell cooperation ^21, 24^. Second, cells export metabolites for many different reasons. For instance, the intracellular biochemical network needs constant balancing which is achieved through an overflow of metabolites ^20, 25–27^. The export of metabolites also mediates metabolic cooperation, a situation that can arise when unrelated biochemical reactions interfere and inhibit each other, and where smaller metabolic networks are more efficient ^28^. Finally, the biosynthetic burden caused by expensive metabolic reactions can be mitigated through the sharing of labour. This helps cells to reduce energetic costs as well as the load of toxic intermediates ^21, 29, 30^.

Intercellular metabolic interactions hence emerge because cells constantly export a broad range of metabolites and, at the same time, harbour an array of mechanisms which sense and uptake extracellular nutrients ^20, 31–33^. This situation has, however, a significant impact on cellular physiology. Once a cell has started to uptake extracellular metabolites, its own biosynthetic pathways typically become inhibited ^32, 34^. As a consequence, metabolic and physiological properties that also impact ageing, such as growth rate, stress tolerance, or the formation or consumption of metabolites such as NADPH, are altered depending on the uptake of metabolites ^32, 35–38^. For example, cells that uptake lysine from the environment mount better protection against oxidants, via increased pools of NADPH ^37^, and cells that rely on the uptake of amino acids, including histidine, leucine, and methionine, are more drug tolerant ^38^.

While there is an increasing understanding of the role of the metabolic environment in stress resilience and growth, the role of cell-cell metabolic interactions in the ageing process is still barely understood ^2^. A measure of eukaryotic cell ageing is chronological lifespan (CLS) in yeast. CLS refers to post-mitotic cell ageing, assessed by the length of time quiescent (non-dividing) cells can survive post entry into the stationary phase ^39, 40^. CLS has been pivotal in the discovery of several of the most critical and conserved regulatory pathways of ageing that are now known to be important across eukaryotes, including the AMPK, mTOR and sirtuin pathways ^8–11^. Monitoring metabolic consumption and production during CLS using metabolomics and isotope tracing experiments, we observe that metabolites exported by young, exponentially growing, cells are re-imported during the stationary phase when cells age chronologically, indicating the existence of cross-generational metabolic interactions. As this result implied that metabolite exchange interactions could be impact CLS, we boosted metabolite export and metabolic interactions through the use of self-establishing metabolically cooperating (SeMeCo) communities, a yeast community model that allows the tracing of metabolite consumer or producer cells of four different metabolic pathways in their sixteen possible combinations (metabotypes) ^41^. We observed a significant extension of the yeast chronological lifespan when cell-cell metabolic interactions are boosted. In the search for the underlying mechanisms, we coupled lifespan assays with proteomics, metabolomics and genome-scale metabolic flux analysis, and discovered a role for the extracellular metabolic environment that is created by the cooperating communities. We find that cells cooperating for the biosynthesis of methionine generate a protective metabolic environment, in which methionine consumers obtain a more glycolytic metabolism and overflow glycolytic products, glycerol in particular. The exometabolome created this way, in turn, extends the lifespan of all cells in the community via a paracrine effect. Our results show that widespread metabolome changes, occurring when cells cooperate metabolically, create a pro-survival metabolic environment leading to extension of their own lifespan. Ultimately, these findings establish cell-cell metabolic interactions and generated exometabolomes as a longevity modulating mechanism.

## Results

### Yeast cells establish cross-generational metabolite exchange interactions during chronologic ageing

As cells sense extracellular metabolites and feedback inhibit their own metabolite synthesis pathways when grown in rich media ^25, 31, 34, 41^, we conducted CLS experiments using synthetic minimal (SM) growth media lacking amino acid and nucleotide supplements. We used a common lab strain in which four artificially introduced metabolic biosynthetic deficiencies (auxotrophies) in three amino acids (*his3Δ1*, *leu2Δ*, *met15Δ*) and one nucleobase (*ura3Δ*) ^42^ were repaired through genomic integration of the wild-type alleles ^38^.

The prototrophic cells, metabolically competent for the biosynthesis of the four metabolites (wild-type cells) were grown in batch culture through exponential, early stationary and stationary phases (1, 2 and 8 days of culture respectively) (**Fig 1a**). Initially, cells grow exponentially (E), consuming glucose supplemented to the culture media, followed by a decline in growth rate as cells transition from diauxic shift to stationary phase (early stationary phase, ES), and then enter stationary phase (S) once preferred carbon sources are exhausted. While exponential cells are mostly glycolytic, they then start consuming released products of glucose catabolism (like ethanol or glycerol) during the diauxic shift, before entering stationary phase, when cells arrest growth and mostly use oxidative phosphorylation to generate ATP (**Fig. 1a**). In order to evaluate the intracellular metabolome of chronologically ageing prototrophic cells, we used a targeted LC-MS/MS method ^43^. The concentration profile of intracellular amino acids, nucleotides as well as glycolysis and tricarboxylic acid (TCA) intermediates was specific to the growth phase; the profiles clustered in a principal component analysis (PCA) according to growth phase (**Fig. 1b i)**). The metabolite concentration changes measured reflected the known metabolic transitions from exponential to the stationary phase ^44–46^. Consistent with a shift from fermentation to oxidative metabolism, the overall concentration of glycolytic metabolites decreased, while we detected an increase in the concentration of TCA derived metabolites (**Fig. 1b ii)**). Moreover, reflecting the ceasing of growth, the concentration of nucleotides decreased during early stationary and stationary phases ^46^. Interestingly, a differentiated picture was obtained for intracellular amino acids. While the concentration of overall amino acids did increase in the stationary phase, we observed a spread in the concentration range (**Fig. 1b ii)**), unpaired two-sided Wilcoxon Rank Sum test and multiple testing correction using the Benjamini & Hochberg (BH) method, adjusted p-values in **Supplementary File 1)**. For example, isoleucine, glycine and leucine increased from exponential to stationary phase, but aspartate, alanine and glutamate decreased, most likely reflecting their role in interconversion reactions in the biosynthesis of other metabolites, including other amino acids and pyruvate (**Fig. 1c**), unpaired two-sided Wilcoxon Rank Sum test and multiple testing correction using the BH method, adjusted p-values in **Supplementary File 2**).

**Figure 1.**
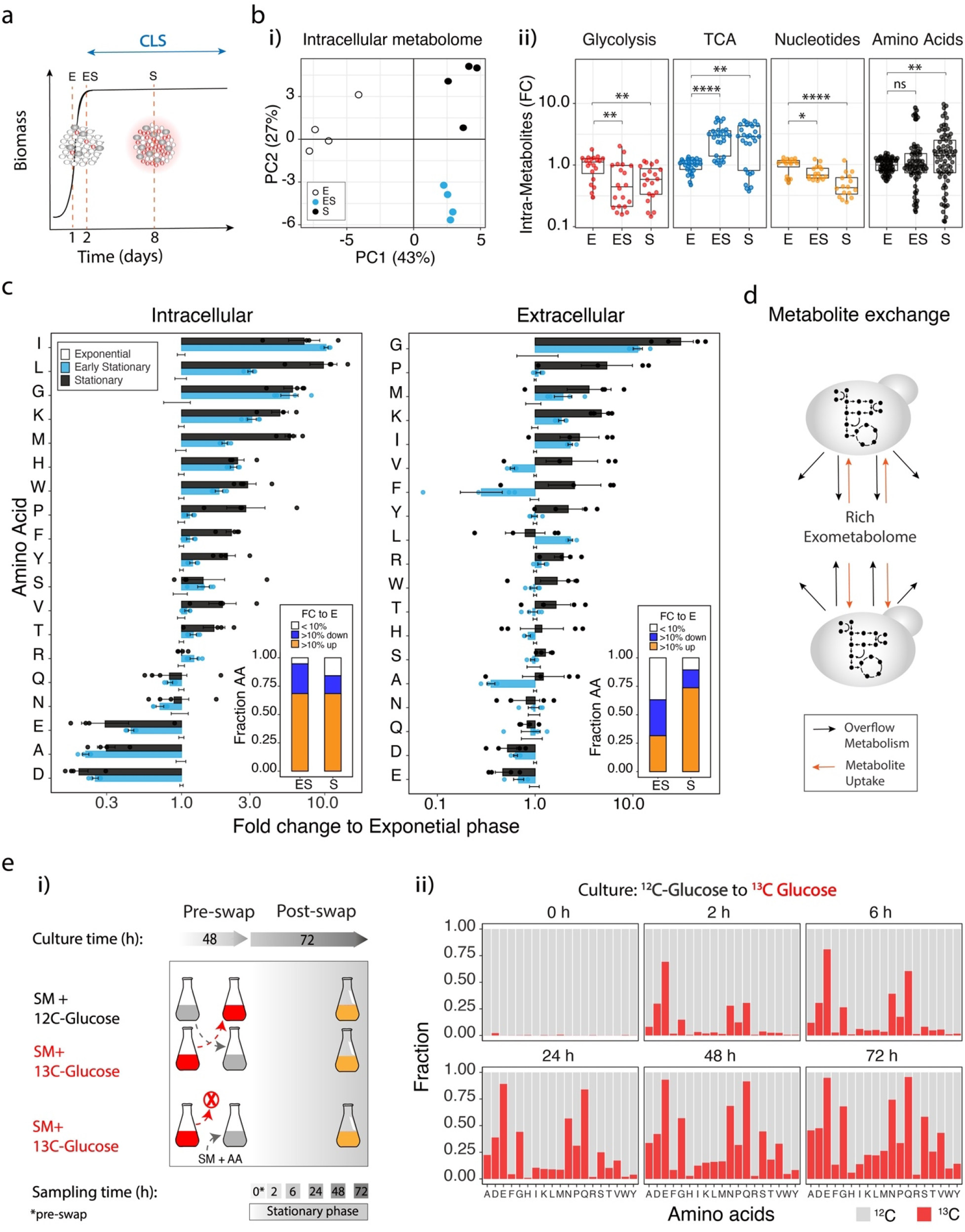
Metabolites exported by exponentially growing yeast cells are imported by chronologically ageing cells in the stationary phase. **a)** Scheme: Growth phases of yeast in batch culture, and the measurement of chronological lifespan (CLS) as survival in the stationary phase. (E) exponential growth phase, (ES) early stationary phase and (S) stationary phase. Red dotted lines indicate the time points of sample collection for metabolomic analysis (1, 2 and 8 days from culture start, indicating E, ES and S phases). b) Targeted intracellular metabolome (intra-metabolome) analysis for the quantification of nucleotides, amino acids, glycolysis and tricarboxylic acid cycle (TCA) metabolites, in wild-type yeast cells cultured in synthetic minimal (SM) medium at 1 (E), 2 (ES) and 8 (S) days from culture start, respectively. i) Principal component analysis (PCA) reflects metabolomic differences between E, ES and S growth phases. Data are n=4 independent cultures (individual dots represent independent cultures). ii) Intracellular metabolite concentrations shown as fold change (FC to exponential phase) of different metabolites according to metabolite classes in the different growth phases. Concentration of each metabolite was first normalised by biomass, as assessed by OD_600_. Box plots represent median (50% quantile, middle line) and lower and upper quantiles (lower (25% quantile) and upper (75% quantile), respectively) of pooled metabolite levels of 4 independent cultures. Each dot represents a metabolite in a biological replicate. Statistics by unpaired two-sided Wilcoxon Rank Sum test and multiple testing correction using the BH method; adjusted p-values *<0.05, **<0.005, ***<0.0005 and ****<0.00005; adjusted p-values are listed in **Supplementary File 1**. c) Intracellular and extracellular amino acids levels in wild-type cultures during exponential (E), early stationary (ES) and stationary (S) phases. Bar plots show mean±SEM fold change (FC to levels in the exponential phase) of n = 4 independent wild-type yeast cultures. Statistics by unpaired two-sided Wilcoxon Rank Sum test and multiple testing correction using the BH method; adjusted p-values are listed in **Supplementary File 2**. Inlets represent the fraction of amino acids (from a total of 19) that show minimal changes, are decreased or increased in the FC to exponential phase (E), as shown by FC<10%, FC >10% down and FC>10% up, respectively. d) Scheme: Cells synthesise metabolites, following the stoichiometric rules of the metabolic network, wherein some metabolites are exported in order to maintain metabolic homeostasis (overflow metabolism), contributing to the metabolic enrichment of the extracellular environment (rich-exometabolome). At the same time, cells can sense and import metabolites from the surrounding environment. These dynamic export/import properties result in the exchange of metabolites between co-growing cells and the establishment of intercellular metabolic interactions. e) i) Scheme: Isotope tracing experimental design. Prototrophic yeast cells were grown in SM media supplemented either with ^12^C-glucose or ^13^C-glucose, during 48 hours, then the media was swapped for isotope tracing amino acid analysis using targeted metabolomics by HPLC-MS/MS ^50^, at 2, 6, 24, 48 and 72 hours post media swap (plus a control 0 h collection, just prior swapping media). Control cultures were swapped from SM supplemented with ^13^C-glucose to SM solely supplemented with ^12^C-amino acids (AA) at standard culturing concentrations (see Methods). ii) Fractions of ^12^C and ^13^C labelled amino acids in cultures initially grown in SM + ^12^C-glucose and swapped to ^13^C-glucose, at different time points post media swap, as described in i). Data represents the mean of 4 independent cultures; Individual fraction values are listed in Supplementary File 3.

As amino acids can be exported by yeast cells into the surrounding environment ^25, 41, 47–49^, we therefore continued our analysis with a quantification of the extracellular amino acid pools using a targeted LC-MS/MS method ^50^. Despite having inoculated our cells in a minimal medium lacking amino acid supplements, we found that by mid-log phase (exponential phase) yeast cells had produced and exported amino acids to reach significant concentrations in the medium. Further, 14 of the 19 analysed amino acids are increased by more than >10% in the extracellular medium in the stationary compared to the exponential phase medium (**Fig. 1c inlet**). Indeed, only glutamate and aspartate were reduced in the extracellular metabolome of stationary cells when compared to the medium formed by exponential cells (**Fig. 1c**, unpaired two-sided Wilcoxon Rank sum test and multiple testing correction with BH method, adjusted p-values in **Supplementary File 2**). The source of these metabolites can be metabolite export as well as cell death in the stationary phase. As most of the metabolites were already increasingly detected in the exponential and early stationary phases where cell death is negligible (∼95% live cells) (**Supplementary Fig. 2a**) and also because in stationary cultures, the concentration of metabolites did not correlate with the number of live cells (**Supplementary Fig. 2b**), we concluded that the main source of metabolites is export during the exponential and early stationary phases.

Amino acids are sensed and efficiently uptaken by actively growing yeast cells ^25, 31, 47^. We therefore asked if cells during the stationary phase, no longer actively proliferating, would uptake the amino acids previously produced in the exponential phase (**Fig. 1d**). We exploited ^13^C-glucose isotope labelling to test for the consumption, by stationary cells, of metabolites that had been produced during the exponential phase. We cultured wild-type yeast cells on SM media supplemented with ^12^C-glucose or ^13^C-glucose for 48h, a duration which ensured that the glucose in the media had been exhausted - catabolized into unlabeled (^12^C) and labelled (^13^C) metabolites, respectively. Then we swapped the media between labelled and unlabelled cells. In parallel, we set control cultures growing on SM media supplemented with ^13^C-glucose, which were then swapped into SM media only supplemented with amino acids (without glucose), to allow distinguishing if intracellular amino acid levels were a direct result of import, or indirectly derived from catabolism of imported carbohydrates. Levels of fully labelled (^13^C) or unlabeled (^12^C) intracellular amino acids (from glucose catabolism) were quantified using a targeted LC-MS/MS method ^50^ (**Fig. 1e i)**). Growth on SM media supplemented with ^12^C-glucose or ^13^C-glucose did not change cell growth parameters prior or post swap (**Supplementary Fig. 3a-b**, unpaired two-sided Wilcoxon Rank sum test; p-values are listed in **Supplementary File 3**). Notably, cells in exponential phase synthesise sufficient amounts of amino acids so that they can be exported and then uptaken by neighbouring cells, as shown by the increased intracellular levels of ^13^C- or ^12^C-containing amino acids in cultures initially grown on ^12^C- or ^13^C-glucose, respectively, or when unlabeled amino acids were added to cells previously cultured in ^13^C-glucose in the control cultures (**Fig. 1e ii), Supplementary Fig. 3c**). Hence amino acids that are produced and exported during the exponential growth phase are taken up by yeast cells during the stationary phase, indicating that yeast cells establish cross-generational metabolite exchange interactions during chronological ageing.

### Metabolite exchange interactions extend lifespan in yeast communities

We next questioned what impact the exchange of metabolites might have on chronological lifespan. The export and import of metabolites cannot be prevented without imposing major metabolic constraints on cells. We overcame this issue by choosing to boost metabolite exchange interactions instead and made use of self-establishing metabolically cooperating communities (SeMeCos) ^41^. SeMeCos exploit the segregation of plasmids that encode for essential metabolic enzymes, to stochastically introduce auxotrophies (metabolic deficiencies), upon which cells can only continue proliferation by exchanging metabolites. Because plasmid segregation continues until a maximum amount of auxotrophic cells is reached, SeMeCos boost metabolite exchange interactions within the communities. Indeed, compared to wild-type cell communities, SeMeCos are characterised by increased metabolite export, an increase in extracellular metabolite concentrations, and increased metabolic interactions ^38^. Despite boosting metabolite exchange interactions, SeMeCos still exploit the native metabolite export and import capacities of yeast cells and do not have artificially altered metabolic pathways or metabolite sensing properties ^38, 41, 51^ (**Fig. 2a**).

**Figure 2.**
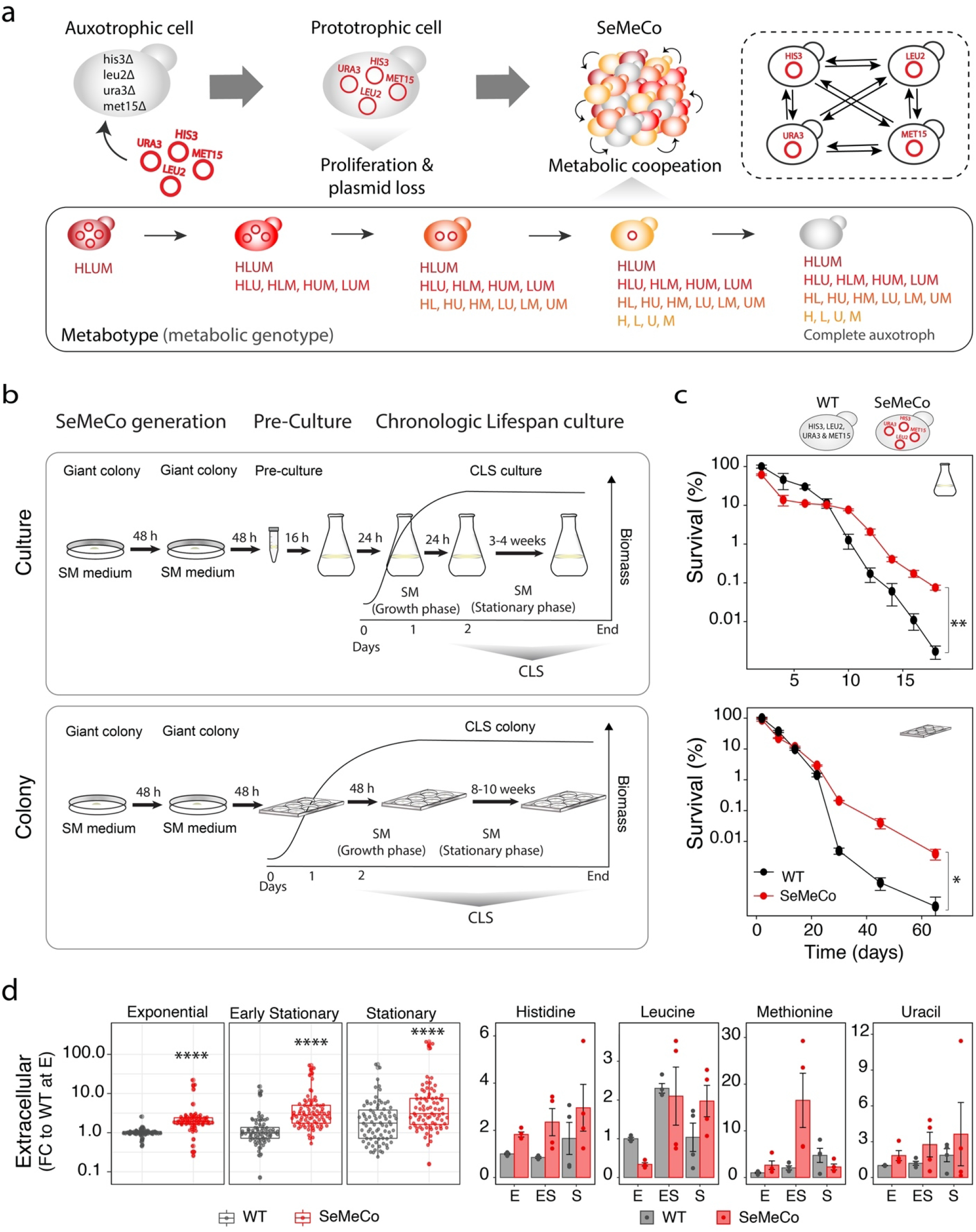
Boosting Metabolite exchange interactions extends chronological lifespan. **a)** Scheme: Self-establishing metabolically cooperating communities (SeMeCos) maximise metabolite exchange interactions through the segregation of plasmid-encoded auxotrophic markers. The SeMeCo system starts with a prototrophic cell that carries four essential metabolic markers on centromeric metastable plasmids (p*HIS3, pLEU2, pMET15* and *pURA3*) instead of stable integration in the genome (prototrophic wild-type). When these cells grow into a community, the stochastic plasmid segregation leads to an increasing number of auxotrophs that continue to proliferate on the basis of metabolites produced by other cells. Segregation continues until a maximum number of auxotrophs that the community can maintain is reached. Sixteen different metabotypes (metabolic genotypes) can arise from the differential segregation of the four metabolic markers. **b)** Chronological lifespan (CLS) of wild-type and SeMeCo communities grown in liquid SM media cultures, start by spotting giant colonies twice to ensure cell proliferation, plasmid segregation and ultimately cross-feeding between auxotrophs and prototrophs (SeMeCos generation), followed by pre-culture and culture, set at high cellular density (assessed by OD_600_) to minimally disturb the composition of SeMeCos in conditions of unsupplemented media. (bottom) CLS of cells grown in a colony follow the same initial SeMeCo generation before re-spotting giant colonies into 6-well plates containing solid SM media. Samples are collected at different time points for survival assessment. **c)** (top) Culture and (bottom) colony CLS analysis assessing survival of wild-type and SeMeCos. Survival was evaluated using colony forming units (CFU) analysis, normalised to biomass (see Methods). Data are mean±SEM survival (percentage fold change) compared to wild-type mean survival at the beginning of stationary phase (48h culture); n = 4 independent cultures per strain (Culture CLS) or n = 3 independent colonies per strain (Colony CLS). Statistics using unpaired two-sided *t*-test, p-value = 0.00661 at day 18 of culture and p-value = 0.0338 at day 65 in the colony; p-values across CLSs are listed in **Supplementary File 4. d)** Extracellular amino acids and uracil levels, measured by HPLC-MS/MS ^50^, in wild-type and SeMeCos cultures during exponential (E), early stationary (ES) and stationary (S) phases. Data are individual metabolite fold-change (FC to mean wild-type levels in the exponential phase) of n = 4 independent cultures per strain. Data comparing wild-type values from Fig 1c; samples were cultured, extracted and measured in parallel. Concentration of each metabolite was first normalised to biomass, as assessed by OD_600_. (left) Box plots showing overall metabolite FC distribution over time; box plots represent median (50% quantile, middle line) and lower and upper quantiles (lower (25% quantile) and upper (75% quantile). (right) Bar plots show the mean±SEM FC of the shared metabolites (HLUM) in SeMeCos over time. Statistics by unpaired two-sided Wilcoxon Rank Sum test and multiple testing correction using the BH method; adjusted p-value *<0.05, **<0.005, ***<0.0005 and ****<0.00005; adjusted p-values are listed in **Supplementary File 6**.

Analysing chronological lifespan of SeMeCos (**Fig. 2b**), we observed that in comparison to the isogenic wild-type strain, SeMeCos were long-lived, as assessed by monitoring colony forming units (CFUs) over time. SeMeCos lost more CFUs immediately after reaching the stationary phase, but in later time-points contained more CFUs and were alive after the wild-type cultures lost viability (**Fig. 2c**, unpaired two-sided *t*-test, p-value = 0.00661 at day 18 of culture; CLS p-values listed in **Supplementary File 4).** To have an independent assessment of survival, we also monitored cell viability using Live/Dead^TM^ cell staining assays. At late timepoints, SeMeCos also contained significantly more viable cells (**Supplementary Fig. 4a**, unpaired two-sided *t*-test; CLS p-values listed in **Supplementary File 4)**. Finally, we exploited the situation where due to the higher cell density and proximity, metabolite exchange is amplified in colonies compared to liquid cultures. Yeast cells survived much longer in colonies than in liquid culture (∼65 vs 20 days). Moreover, also in the colony, SeMeCos had a significantly longer CLS than the isogenic wild-type cells (**Fig. 2c**, unpaired two-sided *t*-test, p-value = 0.0338 at day 65 of growth, CLS p-values listed in **Supplementary File 4)**. We ruled out that the difference in lifespan between SeMeCos and wild-type was explained by differences in pH, a common confounder of lifespan experiments ^52^ (**Supplementary Fig. 4b**, unpaired two-sided *t*-test, p-values listed in **Supplementary File 5).** Moreover, our data suggests that SeMeCo cells were not long-lived due to amino acid starvation, a known lifespan extending intervention ^53^. Indeed, corresponding to the increased exchange of metabolites, the extracellular medium was more metabolite rich in stationary SeMeCos than in wild-type cells, including H, L, M and U **(Fig. 2d, Supplementary Fig. 5**, unpaired two-sided Wilcoxon Rank sum test and multiple testing correction BH method, adjusted p-values in **Supplementary File 6)**.

### The lifespan extension in SeMeCos is mediated by a paracrine mechanism

Next, we tested if the lifespan extension is associated with specific metabotypes, i. e. with specific metabolic dependencies between cells in a community. The SeMeCo model is based on the stochastic segregation of four plasmid-encoded auxotrophic marker enzymes (*HIS3, LEU2, URA3* and *MET15*) resulting in 16 metabotypes (metabolic genotypes, Fig. 2a) distinguishable by genetics. Because of the coupling to other metabolic processes, the 16 metabotypes are connected to broad changes in metabolism and, together, affect differential expression of 2/3rds of the genome ^54^. We monitored occurrence and relative contributions of the different auxotrophies to the ageing SeMeCo cultures over time. Generally, we observed that the proportion of prototrophic cells declined over time and that during the late stationary phase ∼98% of viable cells were auxotrophs. Among the auxotrophs, the *met15Δ* segregants dominated, and increased in relative abundance with time (**Fig. 3a**, paired two-sided *t*-test, p-values in **Supplementary File 7)**.

**Figure 3.**
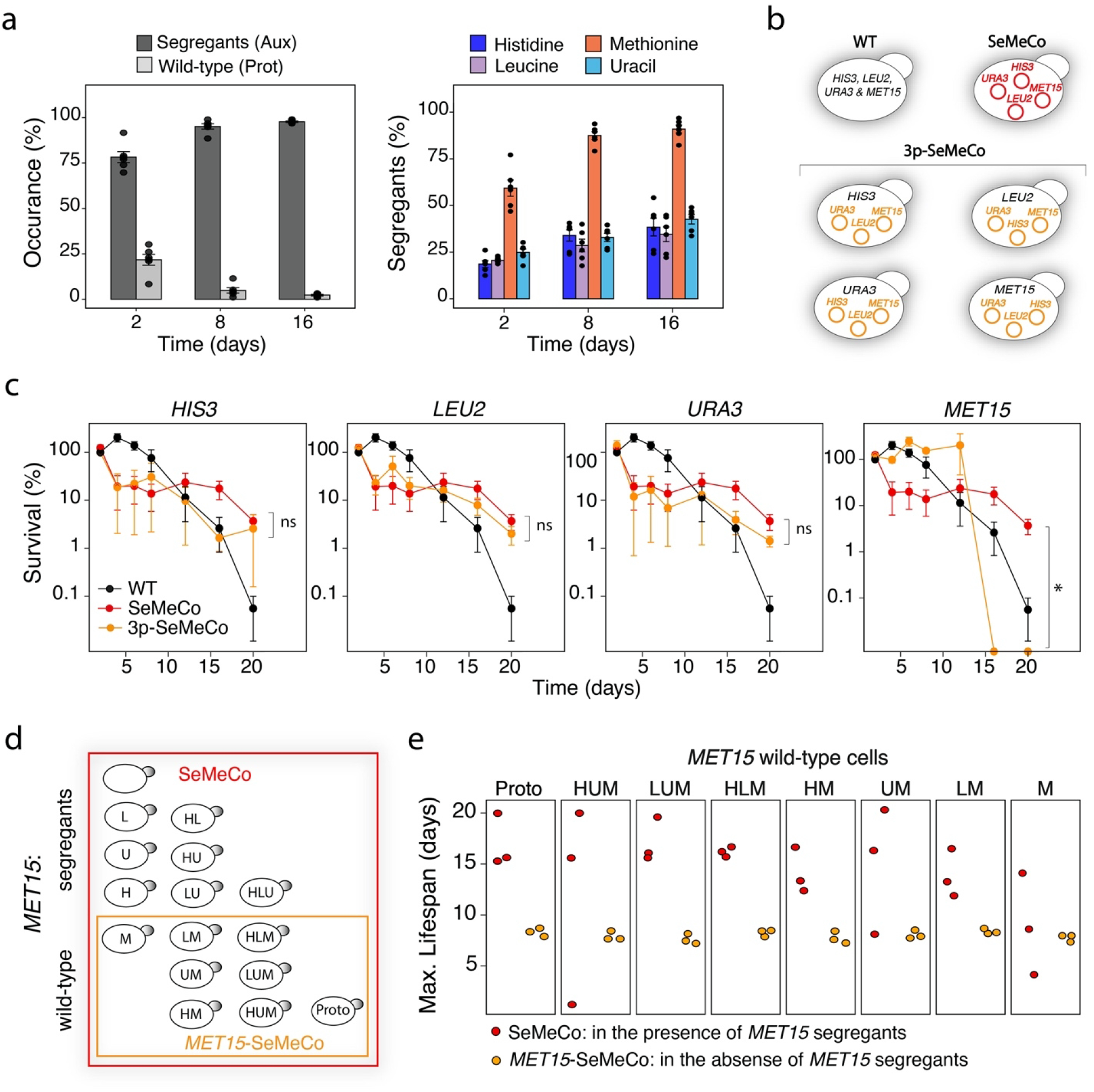
Metabolic interactions between *MET15* and *Δmet15* cells promote a paracrine lifespan extension of SeMeCo communities. **a)** (left) Frequency of overall segregant cells (at least one auxotrophy for H, L, U and/or M) and genetically prototrophic (bearing the four wild-type locus) cells during CLS. (right) Frequency of the individual segregants, i.e. per auxotrophy for H, L, U or M over time. Bar plots show the mean±SEM of n=6 independent SeMeCo cultures across 2 independent experiments (dots represent independent cultures). Statistics by paired two-sided *t*-test; p-values listed in **Supplementary File 7**. The proportion of all auxotrophs increases, with *Δmet15* segregants becoming the most abundant metabotype during CLS. **b)** Schematics: Wild-type, SeMeCos and four ‘3p-SeMeCos’, in which one of the markers (*HIS3, LEU2, MET15 or URA3*) is genomically integrated and does no longer segregate. **c)** Survival percentage of wild-type, SeMeCos and 3p-SeMeCos during CLS (assessed by high-throughput CFU (HTP-CFU) normalised per biomass). Data are mean±SEM survival (percentage fold change) compared to wild-type mean survival at the beginning of stationary phase (48h culture); n=4 independent cultures per strain. Survival curves are shown separately for each 3p-SeMeCos for visualisation purposes (all strains were cultured and analysed in parallel). Statistics by unpaired two-sided Wilcoxon Rank Sum test; p-value at day 20 of culture for: SeMeCo vs wt = 0.0294 SeMeCo vs *HIS3*-SeMeCo = 0.3428, SeMeCo vs *LEU2*-SeMeCo = 0.3428, SeMeCo vs *URA3*-SeMeCo = 0.2000, SeMeCo vs *MET15*-SeMeCo = 0.0210; p-values across CLSs are listed in **Supplementary File 8**. **d)** The segregation of the four metabolic markers gives rise to 16 different metabotypes, eight of which have segregated the *MET15* plasmid (Fig. 2a). **e)** Maximum lifespan of the eight *MET15* wild-type metabotypes, in the presence (SeMeCo, red) or absence (*MET15*-SeMeCo, yellow) of *MET1*5 segregants. Dots are independent cultures per SeMeco type. Data are from n=3 independent cultures per SeMeCo. Statistics by unpaired one-sided Wilcoxon Rank Sum test; p-values in **Supplementary File 11**.

To test the contribution of the individual auxotrophies to the lifespan extension, we next generated additional versions of the SeMeCo communities, in which each one of the auxotrophic markers (*HIS3, LEU2, URA3* or *MET15*) was genomically repaired, and hence, only three plasmids segregate (‘3p-SeMeCos’) (**Fig. 3b**). The genomic repair of *HIS3, LEU2* or *URA3* did not significantly change the lifespan of SeMeCos, whereas the 3p-SeMeco in which the *MET15* locus was no longer segregating had significantly shorter lifespan (**Fig. 3c**, unpaired two-sided Wilcoxon Rank sum test, p-value at day 20 of culture for: SeMeCo vs wt = 0.0294 SeMeCo vs *HIS3*-SeMeCo = 0.3428, SeMeCo vs *LEU2*-SeMeCo = 0.3428, SeMeCo vs *URA3*-SeMeCo = 0.2000, SeMeCo vs *MET15*-SeMeCo = 0.0210; p-values across CLSs are listed in **Supplementary File 8)**. Moreover, SeMeCos that only segregate the *MET15* marker (pM-SeMeCo) also had increased lifespan compared to wild-type communities (**Supplementary Fig. 6**, unpaired two-sided Wilcoxon Rank Sum test, p-value at 28 days of culture = 3.27e-05; p-values across CLS are listed in **Supplementary File 9)**. Both the accumulation of the *met15Δ* segregants during the CLS, and the loss of the phenotype when *MET15* is not segregating, associated the lifespan extension with the methionine biosynthetic pathway and the organic sulphur cycle.

Methionine and other sulphur containing amino acids have repeatedly been linked to ageing, and typically it was a methionine restriction that caused a lifespan extension in model organisms ^55–60^. Interestingly however, the high prevalence of *met15Δ cells in SeMeCos* and the high concentration of methionine produced by cells in the growth media did not suggest that an underlying methionine restriction would apply. To confirm that the fundamental mechanism of our observation is not methionine restriction, we conducted a control experiment, where we supplemented *met15Δ* cells with 2g/L of methionine. Despite the high methionine levels, we observed a robust lifespan extension in *met15Δ* cells (**Supplementary Fig. 6**, unpaired two-sided Wilcoxon Rank Sum test, p-value at 28 days of culture *=* 3.27e-05; p-values across CLS are listed in **Supplementary File 9)**. In search of an alternative explanation, we found evidence for a paracrine effect. We performed an independent CLS experiment, comparing SeMeCos and ‘3p-SeMeCos’ unable to segregate the *MET15* locus, confirmed the dependency of the lifespan extension on the organic sulphur cycle pathway (**Supplementary Fig. 7**, unpaired one-sided Wilcoxon Rank Sum test, p-values in **Supplementary File 10)**, and coupled the CFU analysis with metabotyping analysis (see Methods). Strikingly, we noted that the presence of the *met15Δ* cells significantly increased the maximum lifespan of the other cells in the community (**Fig. 3d-e**, unpaired one-sided Wilcoxon Rank Sum, p-values in **Supplementary File 11)**.

### Lifespan extension in cooperating communities is mediated by an exometabolome rich in protective metabolites

To explore the cell-extrinsic factors that mediate the lifespan extension phenotype, we started by dissecting the metabolic changes emerging when cells metabolically interact. First, we simulated the likely flux changes using a community-adapted version of the flux balance analysis (FBA) that allows monitoring the exchange of metabolites between cells ^38^. We simulated the exchange of metabolites between a prototroph and each of the 15 auxotrophic metabotypes that emerge from all possible combinations of H, L, U and M auxotrophies (**Supplementary Fig. 8a**). The community-FBA revealed that interactions between *MET15* and *met15Δ* cells cause broad metabolic flux changes affecting a range of metabolic pathways, including central metabolism. The broad response involves not only downstream effects of metabolism intracellularly but, in addition to methionine, also results in the exchange of a plethora of other metabolites. (**Supplementary Fig. 8b, Supplementary Fig. 9**, **Supplementary File 12**). This result opened the possibility that it is not only methionine exchange itself, but the metabolic changes introduced by the metabolic cooperation that cause the lifespan extension. In order to get a deeper understanding of the pathways involved, we continued with proteomics analysis. We extracted proteins from the communities, generated tryptic peptides, and analysed them using microLC-SWATH-MS ^61^ and processed the data with DIA-NN ^62^. We then performed differential protein expression analysis comparing otherwise identical SeMeCos that differ in the segregation of the *MET15* marker (SeMeCos vs *MET15*-SeMeCos). We measured proteomes during exponential phase (day 1), early stationary (day 2) and stationary (day 8) growth phases (**Fig. 4a**). We consistently quantified 1951 proteins, around half the typically expressed yeast proteome ^63^. The first principal component (PCA1) in a PCA analysis explained 33% of the variance and separated the proteomes according to growth phase. The second principal component (PCA2), which accounts for a further 23% of the variance, separated the samples based on whether or not the communities segregated the *MET15* marker (SeMeCos vs *MET15*-SeMeCos and wild-type, respectively) (**Fig. 4b, Supplementary Fig. 10**). Moreover, a comparison of the different communities revealed that most of the differential protein expression occurs when *MET15* and *met15Δ* cells interact (**Fig. 4c, Supplementary Fig. 11**, unpaired two-sided *t*-test and multiple testing correction with BH method, adjusted p-values in **Supplementary File 13)**. A Gene Set Enrichment Analysis (GSEA) ^64^ revealed that > 50% of the differentially expressed proteins (unpaired two-sided *t*-test, BH adjusted p-value < of 0.05)) comprised gene ontologies (GO) belonging to metabolic processes (**Supplementary Fig. 12**, **Supplementary File 14)**. Mapping of metabolic enzyme expression levels to the metabolic network allowed visualisation of the changes in metabolism that span over central metabolism and intermediate metabolism, in communities where *MET15* and *met15Δ* cells interact (**Fig. 4d, Supplementary Fig. 13**). Continuing with a pathway-centric analysis of the proteome did point our attention to glycolysis. Enzymes associated with the glycolysis pathway were generally upregulated in the communities that contained *MET15* segregants (**Fig. 4e**). Moreover, an increase in the expression of glycolytic metabolites in stationary cells that typically rely on oxidative phosphorylation for energy production ^44^ was somewhat a surprise (**Fig. 5a-b**). Indeed, both glycolytic activity and glycolytic overflow metabolites are associated with chronological ageing. While glucose restriction itself extends lifespan ^65^, the glycolytic overflow metabolites ethanol and acetate both shorten lifespan ^52, 66^, but another glycolytic overflow metabolite, glycerol, increases CLS ^67^ . We speculated that a change in release of such metabolites might very well change the lifespan of cells that share a common metabolic environment. We therefore measured ethanol, acetate and glycerol in the exometabolome of the different SeMeCos and wild-type communities during CLS. In the stationary phase, levels of all three metabolites were higher in the communities where *MET15* and *met15Δ* segregants interacted. Most striking changes were observed for glycerol, whose levels were ∼8 fold increased, whilst ethanol and acetate levels were ∼2 fold higher (**Fig. 5b i)**, unpaired two-sided *t*-test, p-values in **Supplementary File 15**). In order to explain the sources of the increase in glycerol, we studied the intracellular metabolome. SeMeCos revealed concentration changes in upper and lower glycolytic metabolites across all growth phases: the most significant changes were however detected in the glycerol-associated three carbon phosphates (G3P, DHAP, and PEP) in the stationary phase. These were increased specifically in the communities where *MET15* and *met15Δ* cells interacted (**Fig. 5b ii)**, unpaired two-sided *t*-test, p-values in **Supplementary File 15**). In parallel, we conducted oxygen consumption (OC) analysis. We found that the OC was reduced in the communities containing the *MET15* segregants (**Fig. 5c**, unpaired two-sided Wilcoxon Rank Sum test, p-values in **Supplementary File 16**). The three results were all consistent with the accumulation of glycerol in the extracellular medium: glycolytic enzymes and glycerol precursors were up, while respiratory metabolism, required for the use of a non-fermentable carbon source as glycerol, was reduced. To test whether an accumulation of glycerol could be associated with extending lifespan of cooperating communities, we performed a CLS assay where cells were grown in SM media supplemented with glycerol. Glycerol supplementation extended lifespan to 62 days of culture as compared to previously observed (Fig. 3c) <20 days in wild-type and *MET15*-SeMeCos. The SeMeCos also profited from the glycerol treatment, albeit the relative gain was lower than in wild type cells (mean fold change survival to wild-type in early stationary phase of 5.060%, 0.006% and 0.286% in SeMeCos, wild-type and *MET15*-SeMeCos, respectively, at 62 days of culture) (**Fig. 5d**, unpaired two-sided Wilcoxon Rank Sum test, p-values in **Supplementary File 17)**.

**Figure 4.**
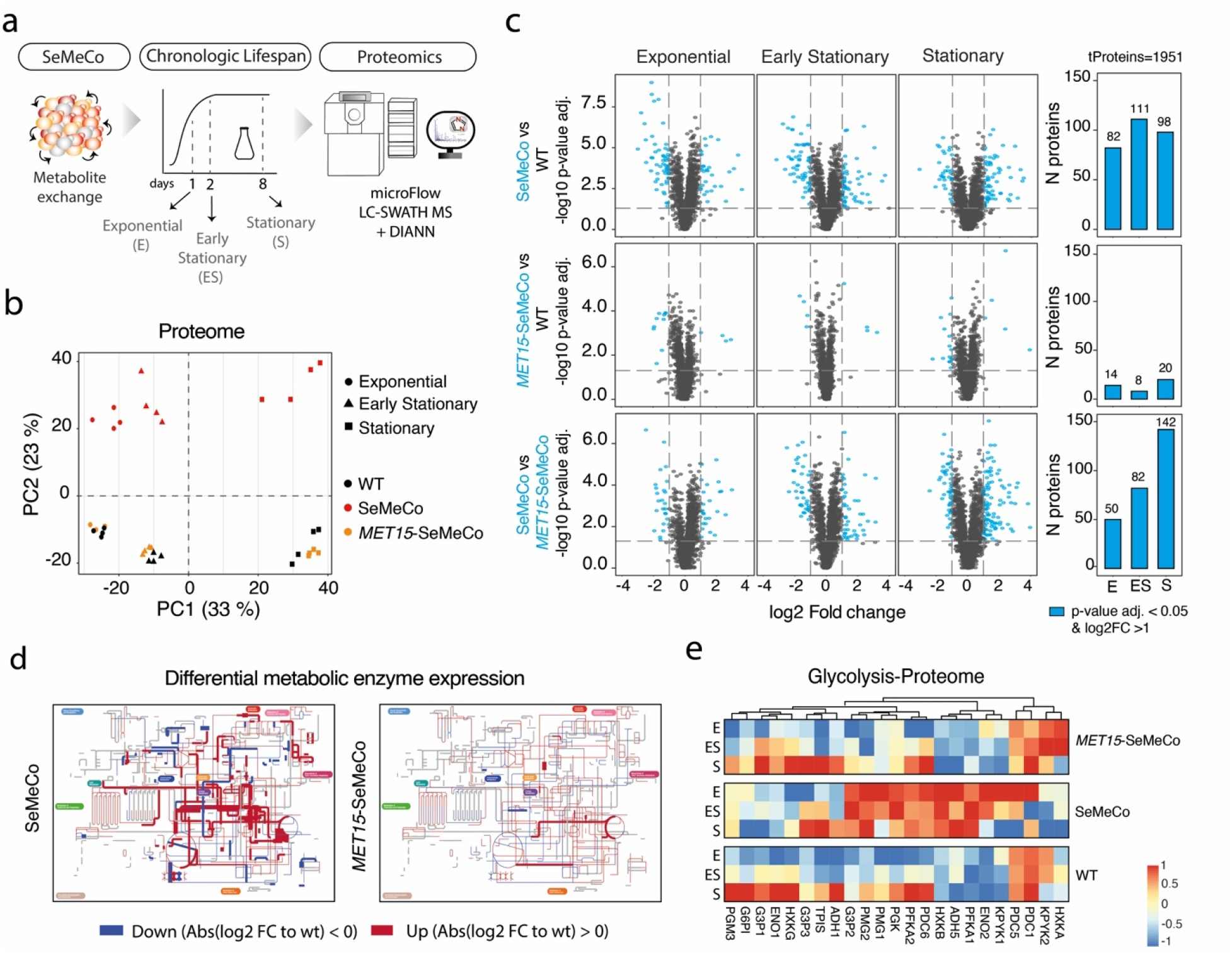
Widespread proteome and metabolome changes in yeast communities where *MET15* and *met15Δ* cells interact. **a)** Wild-type, SeMeCos, and *MET15*-SeMeCos (that do not segregate the *MET15* marker) cells were collected at exponential (E), early stationary (ES) and stationary (S) growth phases. Proteomes were analysed using micro-flow LC-SWATH MS ^61^ and DIA-NN ^62^. Proteomics analysis was performed on four independent cultures (biological replicates) per yeast strain (total n=12 cultures). **b)** Principal component analysis (PCA) reveals that major proteome changes are driven by the transition from exponential to stationary phase (PC1, 33%) and the segregation of the *MET15* marker (PC2, 23%). Individual data points represent biological replicates per strain. **c)** Volcano plots illustrate differential protein expression as log2 fold change (FC) to wild-type expression levels and -log10 adjusted p-value by BH method. Blue dots denote proteins above an absolute log2 FC of 1 (vertical dashed lines) and adjusted p-value <0.05 (horizontal dashed line) (left), with total number of proteins defined as per blue dots represented as bar graphs (right), per growth phase and pairwise comparison. Statistics by unpaired two-sided *t*-test and multiple testing correction using the BH method; adjusted p-values are listed in **Supplementary File 13**. **d)** Differential metabolic enzyme expression levels, from proteome analysis in a), mapped to the yeast metabolic network using IPATH3 ^100^ in the early stationary phase (exponential and stationary phases are shown in Supplemental Fig. 13). Red and blue lines represent significantly (BH adjusted p-value <0.05) up- or down-regulated proteins in SeMeCos and *MET15*-SeMeCos when compared to wild-type; grey lines represent non-mapped/absent proteins in the measured proteomes. Thickness of the lines represent absolute log2 fold change (Abs(log2FC)) changes (thickening = increased Abs(log2FC)). **e)** Expression of enzymes belonging to the glycolysis pathway (columns), derived from the proteome analysis in a) and normalised to a -1 to 1 scale, per growth phase and yeast communities (rows).

**Figure 5.**
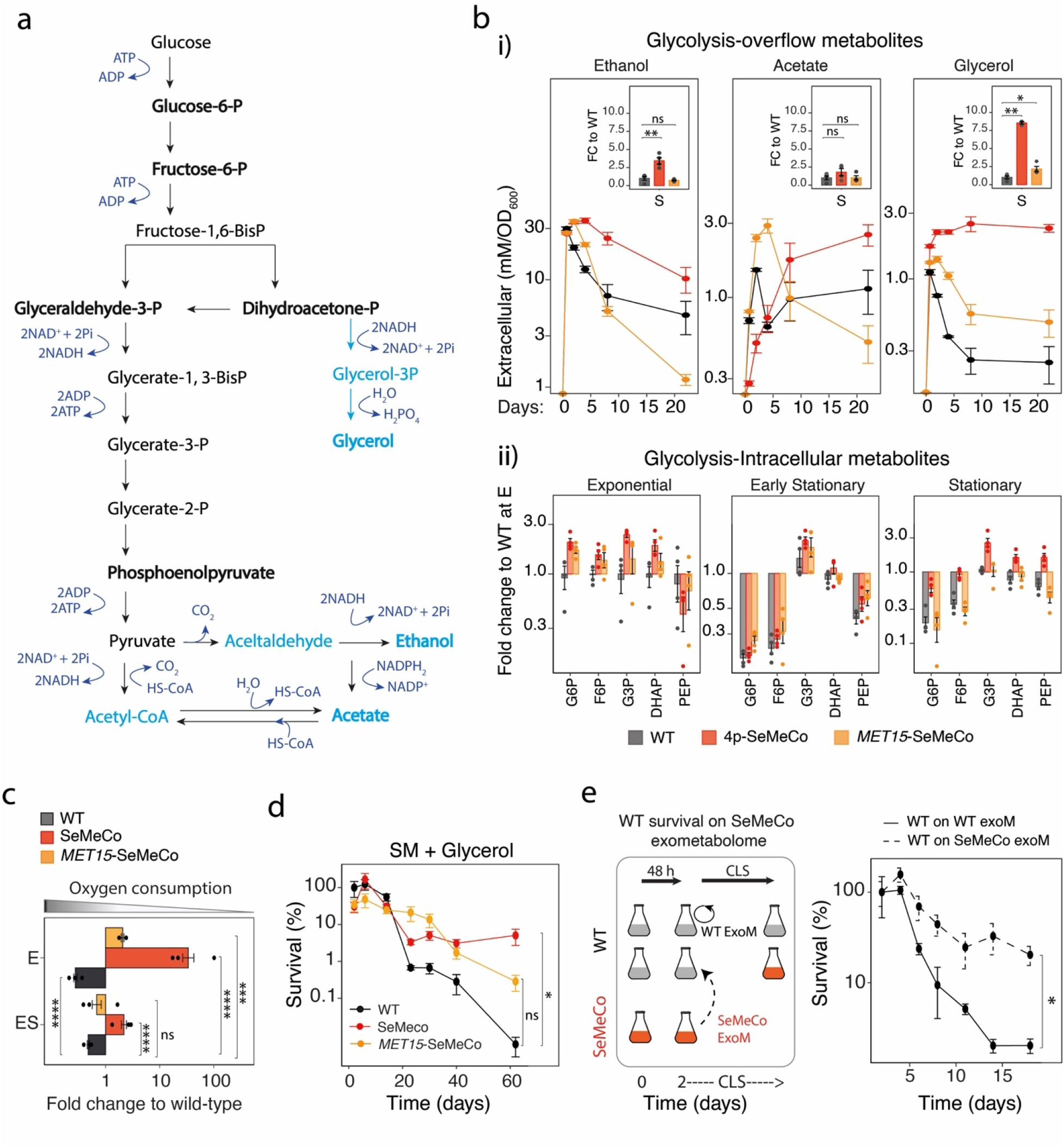
Lifespan extension in SeMeCos is promoted by a self-generated protective exometabolome. **a)** Fermentation (light blue) and glycolysis (black) reaction scheme and associated metabolites **b)** Quantification of **i)** the glycolytic overflow metabolites ethanol, acetate and glycerol in the exometabolome (by HPLC) and **ii)** intracellular glycolytic intermediates (by HPLC-MS/MS) of wild-type, SeMeCos, and *MET15-SeMeCos* that do not segregate the *MET15* gene (top). Data are mean±SEM metabolite levels (mM, normalised to biomass (OD_600_)); n= 4 independent cultures per strain (total n = 12 cultures). Inlets in i) indicate fold change (FC) to mean wild type levels in the stationary phase (S, day 8); individual dots represent independent cultures. Statistics by unpaired two-sided *t*-test; ns= not statistically significant, p-values *<0.05, **<0.005; p-values are listed in **Supplementary File 15**. **c)** Oxygen (O_2_) consumption, as measured by O_2_ saturation in culture post 5 minutes of O_2_ levels recording (using a Hanna HI oxygen meter), in wild-type, SeMeCos, and *MET15-SeMeCos,* that do not segregate the *MET15* gene, cultures in exponential (E) and early stationary (ES) growth phases. O_2_ levels were normalised to biomass, as measured by OD_600_, and to mean levels of wild-type in exponential phase. Data are mean±SEM fold change to wild-type mean levels in stationary phase, n=3 independent cultures per strain (total n=9 cultures). Statistics by unpaired two-sided Wilcoxon Rank Sum test; p-values are listed in **Supplementary File 16**. **d)** Chronologic lifespan assay, shown as survival measured by HTP-CFU normalised to biomass, of wild-type communities, SeMeCos, and *MET15*-SeMeCos, cultured on SM supplemented with glycerol. Data are mean±SEM survival (percentage fold change) compared to mean wild-type survival at the beginning of stationary phase (48h culture); n = 4 independent cultures/ strain (total n=12 cultures). Statistics by unpaired two-sided Wilcoxon Rank Sum test; p-value at day 62 of culture for wt vs SeMeCos = 0.0285 and wt vs *MET15*-SeMeCos = 0.1142, p-values across CLS are listed in **Supplementary File 17**. **e)** Chronologic lifespan assay, shown as survival measured by HTP-CFU normalised to biomass, in wild-type cultures swapped to SeMeCos culture media (SeMeCo exometabolome (exoM)) or kept in their culture media (WT exoM) at the start of stationary phase (48h of growth). Data are mean±SEM survival (percentage fold change) compared to mean wild-type survival at the beginning of stationary phase (48h culture); n=4 independent cultures per exoM. Statistics by unpaired two-sided Wilcoxon Rank Sum test; p-value =0.0286 at day 18 of culture; p-values across CLS are listed in **Supplementary File 18**.

While these results demonstrated that glycerol accumulation is beneficial, the glycerol increase alone might not sufficiently reflect the complexity of the yeast exometabolome. To validate if the community-created exometabolome is indeed mediating the lifespan extension, we hence complemented these results with a media swap experiment. We cultured wild-type communities in SM media until the early stationary phase (48h of culture) and then transferred them to a SeMeCo exometabolome (48h culture media generated in parallel). Control cultures were generated by placing wild-type communities back in their own exomebolome (**Fig. 5e**). Wild-type communities cultured on the exometabolome harvested from SeMeCos showed significant lifespan extension, with mean percentage survival of 20 % in cultures chronologically aged in a SeMeCo exometabolome vs 2% in cultures chronologically aged in a control wild-type exometabolome, representing a 10-fold increase in CFU formation at 18 days of culture, (**Fig. 5e**, unpaired two-sided Wilcoxon Rank Sum test; p-value = 0.0286 at day 18 of culture, p-values across CLS in **Supplementary File 18)**. Hence, the metabolic changes emerging when *MET15 met15Δ* cells interact generate a pro-survival metabolic environment that extends lifespan of all cells in a community.

## Discussion

The classical view of the metabolic network is the one of a biochemical network operating inside the cell. However, with increased understanding of single cell properties, microbial landscapes and phenotypic heterogeneity this view is rapidly evolving ^68^. The individual cell is increasingly seen to be part of a metabolic environment spanning across cells, and thus, metabolite exchange interactions between cells are an essential part of metabolism ^32, 69, 70^. In microbes, metabolic networks span not only over single, but also over multiple species that interact within microbial communities ^36, 49, 71–73^. The degree of metabolite exchange within those communities appears to be extensive. For instance, a majority of microbes are uncultivable outside their community environments, with metabolic co-dependencies being one of the key reasons ^74^. Another interesting observation is that from all 12,538 microbial communities sequenced as part of the Earth Microbiome project ^75^ only 6 contained no amino acid auxotrophs ^38^. The interactions between amino auxotrophs and prototrophs is hence a common situation in microbial communities, which also shows that amino acids are effectively exchanged all the time.

In ecology, metabolite exchange interactions can lead to competition or cooperativity ^21, 23^, but in any case, they have fundamental physiological implications. For instance, we have previously shown that cells that uptake lysine from the environment mount better protection against oxidants ^37^, or that the presence of auxotrophs enriches metabolic environments and increases drug tolerance ^38^. Despite being fundamentally important in modulating cellular processes that also impact on ageing - in particular to growth rate, metabolic signalling, and stress tolerance - to our knowledge, the impact of metabolic intercellular interactions has barely been studied in the context of cellular ageing and lifespan. Indeed, studying the physiological impact of metabolite exchange interactions is technically challenging. Metabolite exchange interactions between cells are not captured by many typical single-cell techniques, such as microscopic imaging or single cells RNA sequencing, nor does the concentration of a metabolite explain whether it was produced or consumed by the analysed cell. Moreover, the export and import of metabolites can not be prevented without imposing major metabolic constraints. We herein hence address this problem by using a combination of various omic, metabolic modelling and genetic techniques, and use synthetic SeMeCo communities to boost metabolite exchange interactions. It is important to stress here that SeMeCos do not rely on new or artificial metabolite exchange interactions to yeast cells; they are based on the exchange of metabolite intermediates, that are a result of the native biosynthetic capacity of yeast, which is boosted through the progressive segregation of plasmids ^41, 51^. Certainly, as a tool of synthetic biology, SeMeCo will not capture all features of natural metabolite exchange interactions. Indeed, with the increasing amount of auxotrophies, SeMeCos become less metabolically versatile compared to prototrophic wild-type cells. However, SeMeCos are one of the very few tools at hand that allow tracing of metabolite exchange between cells, that can flexibly switch between metabolite uptake and consumption. SeMeCos hence prove to be highly useful for studying the physiological consequences of metabolite exchange interactions, by enabling us to boost and trace them.

In studying metabolite exchange interactions in the context of chronological lifespan in yeast, we made two observations that triggered our curiosity. The first was that metabolites exported during the exponential phase, label the cells during the stationary phase, reflecting their import by post-mitotic cells. In the batch-culture, there exists hence a ‘cross-generational’ exchange of metabolites. Ecologically speaking, the batch culture might have been seen as an artificial situation. However, in natural yeast colonies, old and young yeast cells co-exist in close physical distance ^76^. That means that metabolite exchange interactions are even more likely in a natural colony than in batch culture. It is consistent with this notion, that a switch from liquid culture to colony growth tripled the chronological lifespan of our cells. To us, this result hence implies there could be extensive metabolic interactions that apply to both growth and ageing phases of the yeast cell communities.

The second observation was that upon boosting metabolic interactions by using the SeMeCo model, a significant extension of the CLS was achieved. Studying the metabotype composition within SeMeCos attributed a special role to the methionine biosynthetic pathway that is part of the organic sulphur cycle. We observed that cells that had segregated the *MET15* plasmid comprised the highest proportion of long-lived cells in ageing communities. Sulphur amino acids include methionine, cysteine, homocysteine and taurine ^77^ and have previously been associated with lifespan extension. Methionine restriction in particular can extend lifespan in a number of organisms ^55–60^, prevent the development of a variety of diseases ^78^ and influence response to anti-cancer therapies ^79, 80^. Our results differed, however, from many of these studies in a fundamental aspect, as they did not indicate the lifespan extension is caused by methionine restriction. We hence speculated that another mechanism could be at play in the communal cells. The key observation which eventually led to a better understanding of the mechanism, was that the presence of the *MET15* segregants did not only increase their own lifespan, but also the lifespan of the other cells found within SeMeCos. This result strongly suggested that only part of the answer is to be found in the intracellular metabolic reconfiguration in the *MET15* segregants, and that we need to search for a paracrine effect, like a change in the extracellular metabolite pool, to understand the lifespan extension of the entire community.

In order to identify the metabolic changes, we combined metabolite profiling, proteomics and genome-scale metabolic modelling. We detected widespread metabolic changes in the communities containing *MET15* segregants, but were most intrigued by an upregulation of glycolytic enzymes, in a growth phase where typically oxidative metabolism dominates. In following this, we confirmed a decrease in oxygen consumption and an 8-fold increase in the glycolytic overflow metabolite glycerol. Glycerol is known as a protective and pro-survival metabolite ^67^, and also in our hands, significantly extends the lifespan of both wild-type and SeMeCo communities. Glycerol stimulates several survival-associated processes, including osmoregulation, lipid biogenesis, cell wall integrity ^81^ and increase in autophagy ^67^, a known regulator of lifespan ^82^. It is likely that glycerol extends lifespan in a systematic way. Our data does not exclude the possibility that next to glycerol, other protective metabolites enrich in the communal environment, but it shows that in sum, the protective metabolites compensate for the release of acetate and ethanol, that typically result in short-living phenotypes ^52, 66^: the media collected from SeMeCo cells did extend the chronological lifespan of wild-type cells. Our results have interesting evolutionary implications. It has so far been debated if unicellular organisms would profit from a longer lifespan and if they have been selected for longevity ^83^. Our study does not address this question directly, but it reveals an interesting new possibility, in the light that a vast majority of natural microbial communities contain auxotrophs ^38, 74^. Evolution could increase longevity by selecting for populations of cells that metabolically interact in a way that they generate a protective exometabolome. Specifically, the *MET15* segregants, which cannot synthesise methionine and require an organic sulphur source for growth ^84^, cannot survive on their own in the absence of methionine producers. They would benefit if the lifespan of methionine producers is extended. Auxotrophy-prototrohy interactions might hence select for longevity in microbial communities, in other words that metabolic dependencies would not only drive species co-occurrence ^71^ but boost their longevity and evolution. In any case, our results prompt future studies for closely examining the exometabolome as a cause of lifespan extension, specifically when metabolic interventions, such as metabolite restriction/supplementation or metabolic modulating drug treatments are applied.

In summary, we uncover a protective metabolic paracrine effect occurring in metabolically interacting eukaryotic microbial communities. Glycolytic methionine consumer cells enrich the intercellular space for the pro-survival metabolite glycerol, increasing the survival of their producer counterparts and overall community longevity. Impairment or inability to metabolically interact drives cellular dysfunction, which accompanies ageing and disease, therefore dissecting the metabolic dynamics and emerging metabolic environment when cells metabolically interact will aid the development of therapies targeting these processes. Often lifespan extension is associated with restriction conditions, but our data shows that a differentiated view is also necessary, as simple nutritional interventions like the exchange of amino acids can have broad changes in the metabolic network dynamics, reflected in the exometabolome, and alter lifespan this way. Future investigations are necessary to determine how broadly this situation impacts on other nutritional and/or metabolic contexts influencing lifespan.

## Methods

### Yeast cultivation and growth assays

#### Plasmids and strain construction

The haploid BY4741 *S. cerevisiae* strain (*his3Δ1, leu2Δ0, ura3Δ0, met15Δ0*) was used to generate all subsequent strains used in the study. Prototrophy was restored either by genomic knock-in, with primer design based on information from Brachmann *et al.* 1998 ^42^, or plasmid complementation generated by Mülleder *et al.* 2016 ^85^, followed by standard cloning and yeast transformation techniques ^86^. Primers and plasmids details are in Tables S2 and S3, respectively.

#### SeMeCo generation and culture

The generation and culture of SeMeCos was performed as previously described ^38^. The pH, pL, pU and pM plasmid used to generate a SeMeCo strain in the BY4741 background are described in Table S3. All SeMeCo strains were cultured in minimal synthetic (SM) media, composed of yeast nitrogen broth without amino acids (YNB, 6.8 g/L; Sigma #Y0626) + 2% glucose (20 g/L; Sigma #G8270), so cells rely on the exchange of self-synthesised metabolites for growth and survival. Briefly, cryostocks were streaked onto SM + 2% agar medium and cultured at 30 °C for 3 days. Then, a micro-colony was diluted in 500 µl dH_2_O, and normalised to OD_600nm_ = 0.8. Then, 5 µl were spotted onto solid SM medium to generate a giant colony. This initial spotting corresponded to ∼7.2 x 10^4^ cells using a predefined OD-to-cell number standard curve. Cells were incubated for 2 days at 30 °C, then giant colony generation was repeated, to ensure cells have undergone enough proliferation cycles and plasmid segregation, enabling metabolic cooperation, whilst being continuously kept in an exponential growth phase. Pre-cultures were generated by diluting the giant spots into 1 ml dH_2_O, normalised to OD_600nm_ = 1 in SM liquid medium and cultured for 16 hours at 30 °C. Cultures were then generated by diluting the pre-cultures to OD_600nm_ = 0.1 in SM liquid medium and cultured for the duration of the CLS. This relatively high starting OD_600nm_ ensures cells are kept at a density that minimises disturbing the relative proportions of auxotrophs and prototrophs generated in SeMeCos. Cells were collected for downstream experiments at different growth phases, as indicated in figure legends. The control wild-type (BY4741, quadruple knock-in - *HIS3*, *LEU2, URA3 and MET15)* strain followed the exact same procedures as SeMeCos. Strain details are in Table S1.

For CLS assays where cells were grown on glycerol, SM was supplemented with 3% Glycerol (Sigma # G2025) and 1% Glucose; SeMeCo generation and respective wild-type controls were grown on solid SM supplemented with glucose prior to being cultured in SM supplemented with glycerol from pre-culture stage onwards.

#### Knockout strains culture

Knockout strains cultures followed the exact same procedure as described for SeMeCo generation and culture. In the case of metabolic knockout mutants (*met15Δ)*, cells were grown on SM media supplemented with the metabolite for which the strain was biosynthetically impaired (2g/L L-methionine), with respective wild-type controls also being cultured in SM supplemented with the metabolite. Strain details are in Table S1.

#### Isotope tracing

Wild-type yeast cells were cultured in SM media supplemented either with ^12^C-glucose (^12^C-glu; Sigma #G8270) or ^13^C-glucose (^13^C-glu; Sigma #389374), during 48 hours, then media was swapped for tracing amino acid export/import, using targeted metabolomics ^50^ (HPLC-MS/MS), at different time points post media swap. Control cultures were swapped from SM supplemented with ^13^C-glucose to SM solely supplemented with ^12^C-amino acids (^12^C-AA; Sigma #Y1896) at standard culturing concentrations.

#### Exometabolome exchange

Wild-type and SeMeCo yeast cells were cultured in parallel, in SM media, for 48 hours (until the stationary phase). Culture media was then collected by spinning down cells in each culture at 3000g for 5 min at room temperature (RT) followed by supernatant (media) filtering with a 0.22 μm syringe filter. Some wild-type culture cell pellets were then gently resuspended in the filtered media (exometabolome) from SeMeCos whilst others were resuspended back in their own filtered media as control. Wild-type cultures in SeMeCos or wild-type exometabolomes were then followed for CLS.

#### Growth assays

Growth was assessed by monitoring biomass formation using optical density absorbance at a wavelength of 600 nm (OD_600_). OD_600_ was recorded either manually during the CLS, on a Ultrasepc 2100 pro manual^TM^ (Amersham Biosciences), or automatically on a plate reader NanoQuant Plate^TM^, Infinitive 200 PRO (Tecan), every 10 minutes, until cells reached stationary phase, at 30 °C for growth curve recording. Both maximum growth and time to mid log phase were determined from growth curves using the R ‘grothcurver’ package ^87^ .

### Chronologic Lifespan

#### Conventional and High-throughput Colony Forming Unit (CFU) assays

Conventional CFU analysis was performed as described previously ^88^ by aliquoting ageing cultures throughout CLS, and plating cells at different dilution factors into solid rich media (YPD), composed of yeast extract (10 g/L; Sigma), peptone (20 g/L; Sigma), dextrose (20 g/L; Sigma) and agar (20 g/L; Sigma). Cells were incubated for 2 days at 30 °C and the number of CFUs were recorded. Increasing numbers of cultures analysed in parallel required the usage of a high-throughput CFU (HTP-CFU) method as described in ^89^. Briefly, 200 μL aliquots of ageing culture were loaded into the first column of a 96-well plate (8 cultures in parallel per plate). The rest of the plate was loaded with 100 μL of dH_2_O. Using a Biomek NX^P^ automatic liquid handler (Beckman Coulter), 50 μL of the ageing culture from the first column were serially diluted 3-fold across the plate, ensuring each dilution factor was well mixed. Droplets of serially diluted ageing cultures were immediately dispensed onto solid YPD, in quadruplicate (384-well format) using a Singer RoToR HDA pinning robot (Singer Instruments). For this, long-pin 96-density pads were used, making sure that the source plate was revisited before each pin onto the agar. Plates were incubated for 2 days at 30 °C until patterns of colonies appeared. Images of agar plates were acquired with Pyphe-scan ^90^ using an Epson V700 scanner in transmission mode. Plate image analysis and quantification of the number of CFUs in the ageing culture, based on the colony patterns observed, were performed using the R package, DeadOrAlive ^89^. In both conventional- and HTP-CFU assays, survival of the different strains was normalised, first to biomass at time of sample collection, as measured by OD_600_, and then to the survival of the respective wild-type controls at the beginning of the stationary phase (48 hours from the start of culture).

#### Live/Dead Staining

Cell death was assessed using the LIVE/DEAD™ Fixable Far Red Dead Cell Stain Kit, for 633 or 635 nm excitation (ThermoFisher Scientific, Cat no. L10120) according to the manufacturer’s instructions, followed by high-throughput flow cytometry (HTP-FC). Briefly, an aliquot of 300 uL of each ageing culture was transferred to a 96-deep well plate. Plates were then spun down at 3000 g for 3 min RT, supernatant was discarded and cells were resuspended in 300 uL of diluted dye (1:1000 diluted stock dye in dH_2_O), followed by an incubation of 30 minutes in the dark. Cells were then washed in 500 uL of dH_2_O, resuspended in 300 uL of ∼4% formaldehyde (dilute 1:10 37% Formaldehyde in PBS 1x) and incubated 10 min in the dark. Cells were washed in PBS 1x and stored in 500 uL fresh PBS 1x at 4°C, protected from light, until analysis by HTP-FC. Immediately prior to analysis, samples were sonicated for 20 s at 50W (JSP Ultrasonic Cleaner model US21), and 250 uL were transferred to a 96-well plate for HPT-FC analysis. For HTP-FC, 30,000 cells/sample were measured in a Fortessa X20 Flow cytometer (BD Biosciences), using the HTS plate mode on the DIVA software and a 633 nm excitation laser to capture the dye staining. Populations of interest were gated using the FlowJo software version 10.3.0. Features of interest (dead and live cell populations) were then exported for further analysis using R studio.

### Metabotyping

Metabotype performed as previously described ^51^, with the difference that colonies were cryostocked prior to replica-plating, so cells collected at different time points across the CLS would be analysed in parallel. Firstly, conventional CFUs were performed as described above. Then 96 individual CFUs per biological replicate were resuspended in 100 µl of liquid YPD supplemented with 30% glycerol in a 96 well plate (Nunc^TM^ Sigma), as one colony/well, and then incubated at 30 °C ON prior freezing at -80 °C. Once all samples across the CLS were collected, plates were defrosted and then replica plated on six plates, containing either (a) complete medium (SM with all four missing nutrients - 2g/L of L-histine, L-uracil and L-methionine and 6g/L of L-leucine, Sigma), (b) SM medium, and plates with SM and all nutritional supplements except (c) L-histidine, (d) L-leucine, (e) L-uracil, or (f) L-methionine. The absence of growth in a particular drop-out medium reflects the clone auxotrophy for that specific metabolite. The combinatorial growth ability in the six different media allows determination of each clone metabotype (total auxotrophies it contains). This method permits identification of all 16 possible metabotypes resulting from all possible combinations of the four auxotrophies.

### pH analysis

Aliquots of 1 mL were collected per culture at different time points during the CLS and pH was measured using a Mettler-Toledo InLab^®^ Micro & Micro Pro pH electrode coupled to pH/Ion bench meter SC S220-B (Mettler Toledo).

### Oxygen consumption measurements

Ten mL of CLS cultures were collected during exponential (day 1) and early stationary phase (day 2) and placed into a 10-mL Erlenmeyer flask and stirred at 900 rpm using a magnetic stirrer bar. An oxygen probe (Hanna HI 98193), held with a clamp, was inserted into the flask, resulting in it being completely filled with no remaining air inside it, and the flask was sealed with multiple layers of parafilm. The oxygen saturation of the culture was recorded every ∼1 min for 5 min. Oxygen levels were normalised to biomass, as measured by OD_600_, and to levels of wild-type at the end of measurements (5 min).

## Metabolomics

### Targeted metabolomics for intracellular glycolytic and TCA intermediates, nucleotides and amino acids

#### Sample preparation

Ageing cultures, at several time points reflecting different growth phases, were sampled and 400 uL of each culture were quenched in 1600 uL dry-ice-cold methanol, into a 48-deep-well plate. This suspension was spun down (600 g, 3 min, 4°C), and the supernatant was discarded by inversion, followed by a short spin (600 g, 1 min, 4°C) to ensure complete removal of the SN. Cell pellets were immediately placed on dry ice and then transferred to −80 °C until analysis. Intracellular metabolites were then extracted as described ^91^. Briefly, 140 μl of 10:4 MeOH/water were added and vortexed. Then, 50 μl chloroform was added, followed by 50 μl water and 50 μl chloroform with thorough mixing in between. Phases were separated by centrifugation at 3,000 *g* for 10 min. The aqueous phase was recovered and used without further conditioning. One microlitre was injected for HPLC-MS/MS analysis. Before analysis by HPLC-MS/MS the order of samples was randomised and during analysis a quality control sample (QC) was assessed every 24 samples.

#### Sample acquisition

Metabolites were resolved on an Agilent 1290 liquid chromatography system by HILIC coupled to an Agilent 6470 triple quadrupole instrument operating in dynamic MRM mode, as previously described ^43^. In short, the gradient program started at 30% B (100 mM ammonium carbonate) and was kept constant for 3 min before a steady increase to 60% B over 4 min. Solvent B was maintained at 60% for 1 min before returning to initial conditions. The column was washed and equilibrated for 2 min resulting in a total analysis time of 10 min. We used acetonitrile as solvent A and a Waters BEH Amide column (2.1 × 100 mm, 1.7 μm particle size) for separation. The flow rate was set to 0.3 ml/min and column temperature to 35C. Compounds were identified by comparing retention time and fragmentation patterns with analytical standards. Metabolite quantifications were then normalised per biomass, as measured by OD_600_, at the time of collection.

### Targeted metabolomics for intracellular and extracellular amino acids & uracil quantification

#### Sample preparation

Ageing cultures, at several time points reflecting different growth phases, were sampled and 500 uL of each culture were collected into a 96-deep-well plate for amino acid & uracil profiling. Samples were centrifuged at 4,000 g for 3 min and supernatants (SN) were transferred into a new 96-deep-well plate for extracellular metabolite profiling, whilst cell pellets were washed once in dH_2_O, spun down at 4,000 g for 3 min and SN was discarded (followed by a 1 min spin for complete removal of SN) for later intracellular metabolite profiling. Both cell pellets and SN were immediately frozen in dry ice and samples were then stored at −80 °C until analysis.

The amino acid extraction and uracil extraction, separation and detection protocols were adapted from ^50^. Briefly, 200 µl of 80% ethanol at 80 °C were added to the yeast pellets. Samples were heated for 2 min at 80 °C, vigorously mixed on a vortex mixer and incubated for further 2 min at 80 °C followed by vigorous vortexing. The extracts were removed from debris by centrifugation at 12,000 *g* for 5 min. SN were also centrifuged at 12,000 *g* for 5 min to further purify samples from any debris. Before analysis by HPLC-MS/MS the order of samples was randomised and during analysis a quality control sample (QC) was assessed every 24 samples.

#### Sample acquisition

Analysis by LC-MS/MS, amino acids & uracil were separated by hydrophilic interaction liquid chromatography (HILIC) using an ACQUITY UPLC BEH amide column (130Å, 1.7 mm, 2.1 mm X 100 mm) on a liquid chromatography (Agilent 1290 Infinity) and tandem mass spectrometry (Agilent 6460) system. Buffer A was composed of 50:50 acetonitrile/water (Greyhound Bio-012041, Greyhound 23214125), 10 mM ammonium formate (Fluka, Cat. No. 14266), 0.176% formic acid (Fluka, Cat. No. O6454) and buffer B of 95:5:5 acetonitrile/methanol/water (Greyhound BIO-13684102), 10 mM ammonium formate, 0.176% formic acid. The gradient elution was performed at a constant flow rate of 0.9 ml/min. Starting conditions were 85% buffer B, after 0.7 min the concentration of buffer B was decreased gradually to 5% until 2.55 min and kept for a further 0.05 min before returning to initial conditions. The column was then equilibrated, resulting in a total runtime of 3.25 min. Compounds were identified by matching retention time and fragmentation (MS2) with commercially obtained standards (Sigma-Aldrich, Cat. No. LAA21). Signals for free amino acids were then acquired in dynamic SRM mode in the MassHunter Software Agilent. Amino acid & uracil quantifications were then normalised per biomass, as measured by OD_600_, at the time of collection.

Extracellular amino acids and uracil data from wild-type in exponential phase are a re-analysis of data in ^38^; experiments, including cell culture, metabolite extraction and sample acquisition, were performed in parallel.

### HPLC method for ethanol, acetate and glycerol exometabolome quantification

#### Sample preparation

Frozen SN in 96 deep-well plates (collected as described above for amino acid and uracil analysis) were defrosted and kept shaking using plate shaker for 20 minutes 900 rpm room temperature, just before the filtration, using a multiscreen filtered plate with 0.45 um durapore membrane (MVHVN4525) and Strata well plate manifold (https://phenomenex.blob.core.windows.net/documents/863d86a0-3aba-4591-979b-bf54b1188038.pdf) and a Welch vacuum pump.

#### Sample acquisition

The target compounds were quantified using a Shimadzou Prominance HPLC (https://www.ssi.shimadzu.com/products/liquid-chromatography/prominence-hplc.html) equipped with a refractive index detector RID20A and a Sil20-ACT auto sampler with a 96 well plate injector tray. The separation was performed on an Agilent Hi-plex H column (https://www.agilent.com/cs/library/applications/5990-8801EN%20Hi-Plex %20Compendium.pdf). The temperature of the column and detector was 50 and 41 °C, respectively. The eluent was 0.00125 N H2SO4 in Type 1 water (0.6 mL min−1). The samples were kept in 96 well plates (https://www.sarstedt.com/en/products/laboratory/cell-tissue-culture/cultivation/product/83.3926/) covered with silicone mat (https://www.phenomenex.com/Products/Part/AH0-8633) at 4 °C in the autosampler prior the injection for no longer than 2 days. 5 uL was injected from the samples / well plates as well as standard compounds. The method works with 26 minutes cycle time. To keep the retention times and detector response constant 5 L of eluent was mixed in one batch.

For the data analysis the Shimadzou data processing software was used. Target compounds were identified by using automatic retention time matching with individual standards of in the house overflow metabolite library dissolved in SD minimal media. Compound concentrations were calculated using peak area integration with pre-optimized integration parameters and external calibration for each compound. All the calibration curves showed high linearity R2 > 0.9999 at 3 orders of magnitude concertation range. The integration and compound identification were manually overviewed. Data was then exported and further processed using R. Metabolite quantifications were then normalised per biomass, as assessed by OD_600_, at the time of collection.

### Proteomics

#### Sample preparation

Ageing cultures, at several time points reflecting different growth phases, were sampled and 500 uL of each culture were collected into a 96-deep-well plate. Samples were centrifuged at 4,000 g for 3 min and supernatants (SN) were discarded. Samples were centrifuged again at 4,000 g for 1 min to fully remove any residual SN. Cell pellets were immediately placed on dry ice before being stored at -80°C, until all samples were collected. Sample preparation for proteomics was performed as previously described ^61^. Briefly, cell pellets were processed in a bead beater for 5 min at 1,500 r.p.m. (Spex Geno/Grinder), in a lysis buffer where proteins were denatured in 8 M urea (Sigma-Aldrich, 33247) plus 0.1 M ammonium bicarbonate (Sigma-Aldrich, 09830) at pH 8.0. Samples were spun down for 1 min at 4000 r.p.m, before they were reduced in 5 mM dithiothreitol (Sigma-Aldrich, 43815) for 1h at 30 °C and then alkylated in 10 mM iodoacetamide (Sigma-Aldrich, I1149) for 30 min at RT protected from light. Samples were diluted to less than 1.5 M urea in 0.1 M ammonium bicarbonate at pH 8.0, before proteins were digested overnight at 37 °C with trypsin (Promega, V511X). Trypsin was neutralised with 1% formic acid (FA) (Fisher Scientific, 13454279), before peptides were cleaned-up in 96-well MacroSpin plates (Nest Group): 1. plates were first equilibrated in a series of Methanol (1x) (Greyhound Chromatography, BIO-13684102), 50% ACN (2x) (Greyhound Chromatography, Bio-012041-2.5L), and 3% ACN 0.1% FA (2x), between each wash plates were spun down for 1 min at 100 g and flow through was discarded; 2. samples were loaded into the plates and peptides were cleaned-up in a series of 3% ACN, 0.1% FA (3x), between each wash samples were spun down for 1 min at 100 x g and flow through was discarded; 3. peptides were eluted into a new collection plate with 50% ACN (3x) and spun dried overnight at RT in speed vacuum. Peptides were then dissolved in 40 uL of 3% ACN 0.1% FA. Peptide concentration was measured at Absorbance 280 nm using the Lunatic (Unchained Labs).

#### Sample acquisition

The digested peptides were analysed on a nanoAcquity (Waters) (running as 5 µl min^−1^ microflow liquid chromatography) coupled to a TripleTOF 6600 (SCIEX). Protein digest (2 µg) was injected and the peptides were separated with a 23 min non-linear gradient starting with 4% acetonitrile in 0.1 % formic acid and increasing to 36% acetonitrile in 0.1% formic acid. A Waters HSS T3 column (150 mm × 300 µm, 1.8 µm particles) was used. The DIA method consisted of an MS1 scan from m/z 400 to m/z 1250 (50 ms accumulation time) and 40 MS2 scans (35 ms accumulation time) with a variable precursor isolation width covering the mass range from m/z 400 to m/z 1250. Data quantification was performed using DIA-NN version 1.7.1 software ^62^. Post-processing data analysis was conducted in R ^92^.

### Genome-scale metabolic modelling (Flux balance analysis)

#### Constructing auxotroph-prototroph community metabolic models

The community metabolic models were reconstructed using the approach from our previous study ^38^. Briefly, the genome-scale metabolic model of S. cerevisiae ^93, 94^ was used to create auxotrophic strain models by switching off respective metabolic reactions. Then the reactions from auxotroph (H, L, U and/or M) and prototroph (WT) models were combined to make the community, using the compartment per guild approach, where both strains were treated as separate compartments and metabolic exchange between strains were allowed. The community biomass was the combined biomass of all strains. The Cobra toolbox ^95^ was used to perform the model simulations.

### Data analysis and statistics

All statistical analyses were done in R (R Core Team, 2015) ^92^ using specific packages as indicated throughout the methods section. For the basic data manipulation and visualisation we used the R tidyverse package compilation and for statistical analysis we used the R ggpubr package. Hypothesis testing to assess means of population differences were mainly done using *t*-test whenever the variables could be assumed continuous, or otherwise using Wilcoxon Rank Sum test, as indicated in the respective figure legends. Sample size estimations were not performed in any of the experiments. All experiments were performed using at least n=3 biological replicates. Post-processing data analysis was conducted in R. Missing values in the proteomics data were median imputed. Differential protein expression analysis was performed using the limma package v3.48.3 in R ^96^. Gene Ontology (GO) terms were retrieved using GO2ALLORFS object of org.Sc.sgd.db v3.14.0 package ^97^ and enrichment analysis of differentially expressed proteins was performed using hypergeometric statistical tests. GO slim term mapper from SGD database ^98^ was used to map differentially expressed proteins with GO slim terms. KEGG term mapper from KEGG database ^99^ used to map differentially expressed proteins with KEGG terms. Metabolic enzyme expression levels were mapped to the yeast metabolic network using iPATH3 ^100^.

### Data availability

The data supporting the findings of this study are available within the paper, its Supplementary Information and will be deposited within publicly accessible repositories (before formal acceptance). The proteomic datasets generated during the current study that are relevant to data shown in Fig. 4 and Supplementary Fig 10-13 will be available from the PRoteomics IDEntifications database (PRIDE, https://www.ebi.ac.uk/pride/) as part of the global Proteomexchange (PX) consortium ^101^. Yeast gene functions and GO slim term mapper can be accessed at the Saccharomyces Genome Database (SGD, https://www.yeastgenome.org/). Protein sequence databases used for the identification and mapping of proteins from proteomics can be accessed via Uniprot (https://www.uniprot.org/) and KEGG (https://www.genome.jp/kegg/pathway.html), respectively.

### Code availability

No custom codes were generated as part of this study. All analyses conducted in R v3.6.1 used standard, publicly accessible packages obtained either through GitHub (https://github.com/), the Comprehensive R Archive Network (CRAN, https://cran.r-project.org/) or via Bioconductor (https://www.bioconductor.org/).

## Supporting information

Supplemental Files for statistical data

## Acknowledgements

We thank our lab member Dr. Benjamin Heineike (The Francis Crick Institute, London, UK) for inspiring discussions. This work was supported by the Francis Crick Institute, which receives its core funding from Cancer Research UK (FC001134), the UK Medical Research Council (FC001134) and the Wellcome Trust (FC001134, IA 200829/Z/16/Z). This research was funded in part by the Wellcome Trust (FC001134 and IA 200829/Z/16/Z to M.R., supporting C.C-M., S.K., M.M. C.B.M., L.H-D., V.D., SJ.T., A.F., K.C., S.A., L.S. and J.S.L.Y). For the purpose of Open Access, the author has applied a CC BY public copyright licence to any Author Accepted Manuscript version arising from this submission. The work was further supported by the Ministry of Education and Research (BMBF), as part of the National Research Node ‘Mass spectrometry in Systems Medicine (MSCoreSys)’, under grant agreements 031L0220 (to M.R.) and 161L0221 (to V.D.), and the European Commission (EC) as part of CoBioTech project Sycolim ID#33. M.T.A. is funded by the United Arab Emirates University, Al Ain (UAE). The work performed by R.T. is supported by the National Research, Development and Innovation Office PD 128271 and B.P. is supported by the National Research, Development and Innovation Office Élvonal Program KKP 129814, Lendület” program of the Hungarian Academy of Sciences LP2009-013/2012 and the European Union’s Horizon 2020 research and innovation program Grant No. 739593. AZ was supported by the SciLifeLab funding and Marius Jason Jakulis (MJJ) foundation.

## Author contributions

Conceptualization was carried out by C.C-M. and M.R. Methodology was the responsibility of C.C-M., S.K., M.M., C.B.M, SJ.T and L.S. Experimental investigation and experimental design was performed by C.C-M., S.K., M.M., L.H-D., C.B.M. R.T., A.F., K.C., B.P., S.A., L.S and J.S.L.Y) Formal analysis was conducted by C.C-M., V.D., A.Z., SJ.T., and M.T.A. C.C-M and M.R. wrote the article.

## List of Supplementary Information

● Supplementary Figures: 1-13
● Supplementary Tables:1-3
● Supplementary Files: 1-18 (provided as .xlxs or.csv)

