## Supplemental_Information for "Cell-cell metabolite exchange creates a pro-survival metabolic environment that extends lifespan"

**List of Supplementary Information:**

Supplementary Figures: 13

Supplementary Tables: 3

Supplementary Files: 18 (provided as .xlxs or.csv)

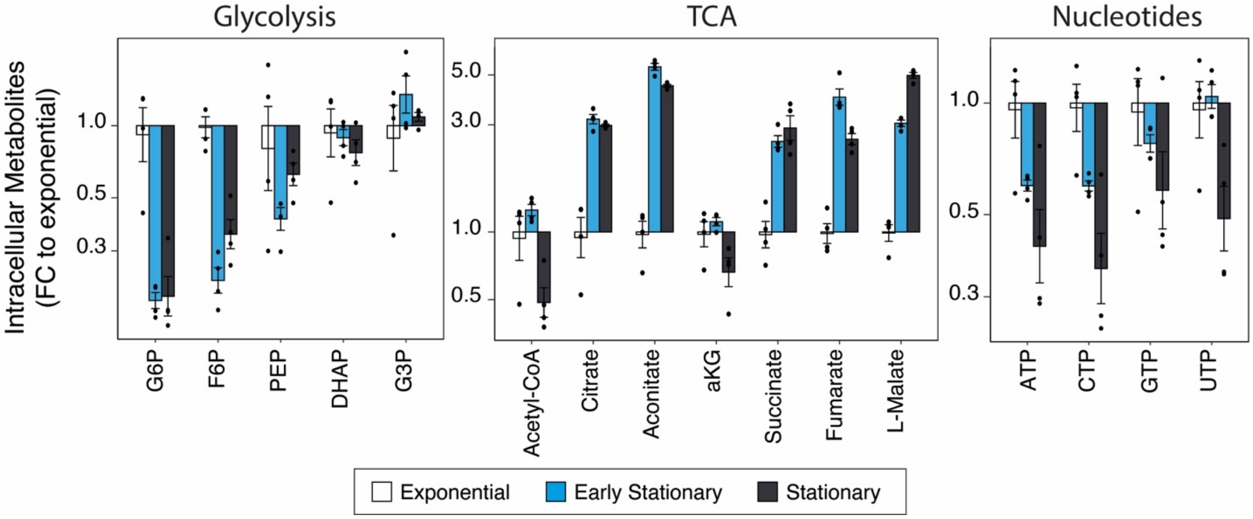


**Supplementary Fig. 1 Intracellular metabolites levels during chronologic ageing. a)** Quantification of intracellular metabolites levels by targeted metabolomics (HPLC-MS/MS), categorised by metabolic pathway, during exponential, early stationary and stationary growth phases. Bar plots show mean±SEM fold change (FC) to levels in the exponential phase of n= 4 independent wild-type yeast cultures; individual dots represent independent cultures. Statistics by unpaired two-sided Wilcoxon Rank Sum test and multiple testing correction using the BH method; adjusted p-values are listed in **Supplementary File 1**.

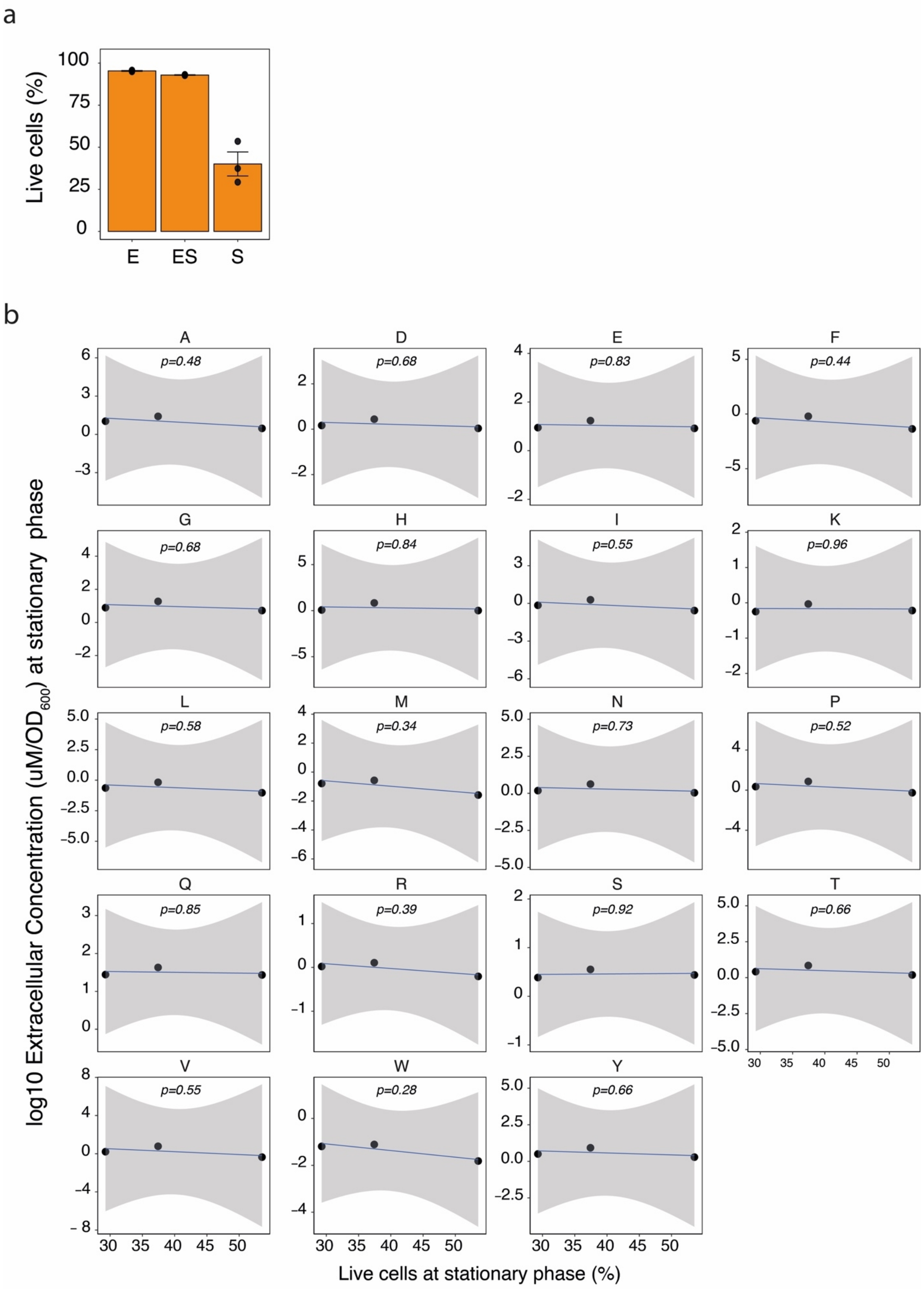
a
Live cells (%)
100
75
50
25
0
E
ES
S
b
A
p=0.48
D
p=0.68
E
p=0.83
F
p=0.44
G
p=0.68
H
p=0.84
I
p=0.55
K
p=0.96
L
p=0.58
M
p=0.34
N
p=0.73
P
p=0.52
Q
p=0.85
R
p=0.39
S
p=0.92
T
p=0.66
V
p=0.55
W
p=0.28
Y
p=0.66
log10 Extracellular Concentration (uM/OD600) at stationary phase
Live cells at stationary phase (%)

**Supplementary Fig. 2 Cell death does not define the increased extracellular amino acid levels during chronologic ageing. a)** Wild-type yeast cultures grown in SM media and collected in exponential (E), early stationary (ES) and stationary (S) phases for cell viability assays, using the LIVE/DEAD™ fixable dye. Bars plots show the percentage of live cells in the different yeast culture growth phases. Data are mean±SEM of n = 3 independent cultures (dots represent independent cultures). **b)** Pearson correlation between extracellular amino acid levels (normalised by biomass, as assessed by $OD_{600,}$ from Figure 1c) and the percentage of live cells, as measured by staining with LIVE/DEAD™ fixable dye, in stationary phase cultures (day 8 of culture). Data are from n = 3 independent cultures (dots represent individual cultures); error bands indicate the 95% confidence level interval for the predictions from the linear model.

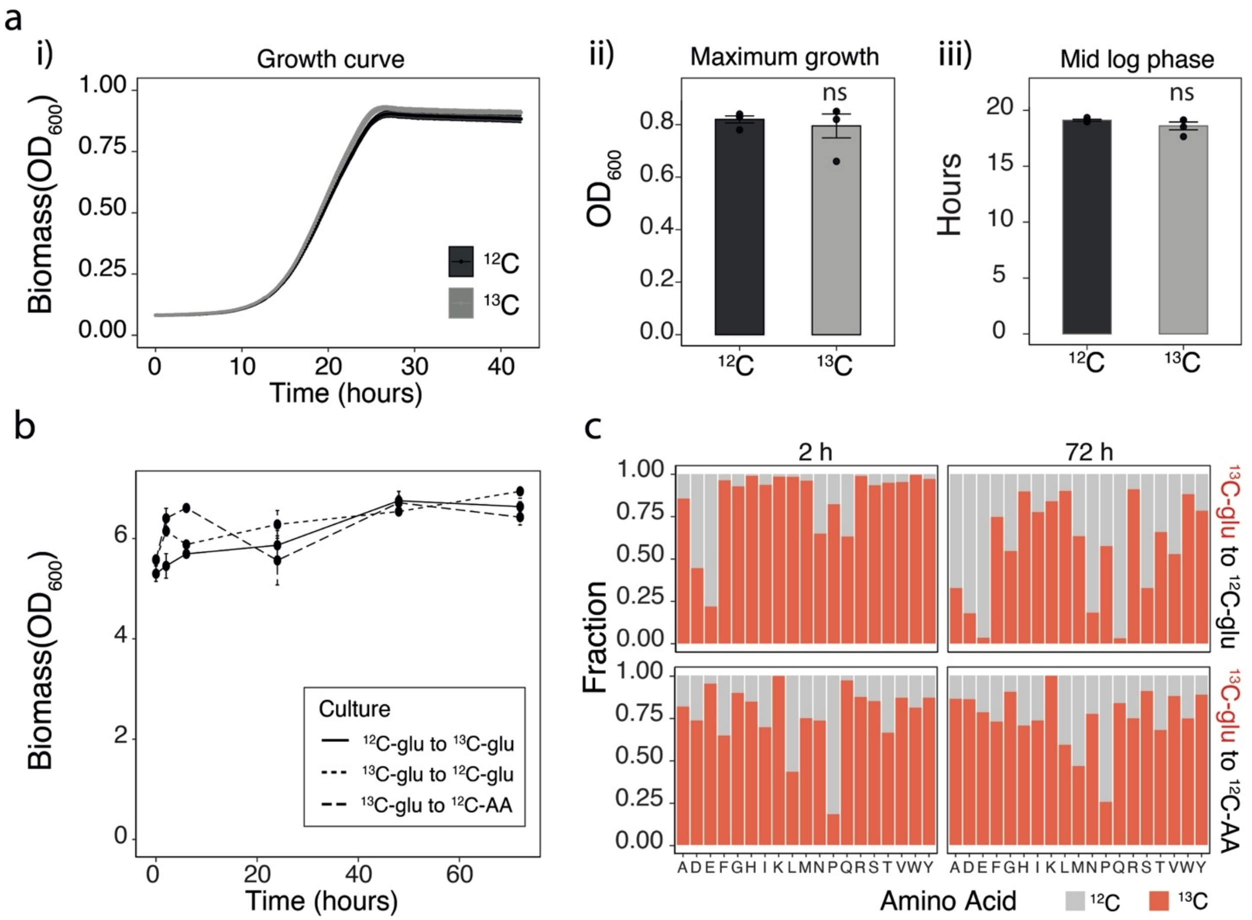


**Supplementary Fig. 3 High degree of import of secreted amino acids by cells during stationary phase.** Wild-type yeast cells were cultured in SM media supplemented either with $^{12}$C-glucose ($^{12}$C-glu) or $^{13}$C-glucose ($^{13}$C-glu) for 48 hours, then media was swapped for tracing amino acid export/import, using targeted metabolomics (HPLC-MS/MS), at different time points post media swap. Control cultures were swapped from SM supplemented with $^{13}$C-glucose to SM solely supplemented with $^{12}$C-amino acids ($^{12}$C-AA) at standard culturing concentrations. **a)** Prior media swap growth curves (i), maximum growth (ii) and time to mid log phase (iii), as measured by $OD_{600}$, of wild-type yeast cells grown on SM media supplemented either with 2% $^{12}$C-glucose ($^{12}$C-glu) or $^{13}$C-glucose ($^{13}$C-glu). Data mean±SEM of n= 4 independent cultures/per condition (culture media), n= 8 cultures in total. Statistics by unpaired two-sided Wilcoxon Rank Sum test; p-value = ns (not statistically significant); maximum growth p-value=0.883; time to mid log phase p-value=0.686. **b)** Biomass, as assessed by $OD_{600}$, measured at 0, 2, 6, 24, 48 and 72 hours, post media swap from SM + $^{13}$C-glucose to $^{12}$C-glucose ($^{13}$C-glu to $^{12}$C-glu), or SM +

$^{12}C$-amino acids ($^{13}C$-glu to $^{12}C$-AA), or from SM + $^{12}C$-glucose to SM + $^{13}C$-glucose ($^{12}C$-glu to $^{13}C$-glu). The gradual slow increase in biomass, from 0 to 72 hours, in the $^{12}C$-glu to $^{13}C$-glu and $^{13}C$-glu to $^{12}C$-glu exhausted (post 48h) cultures, most likely reflect consumption of preferred carbon sources from overflow glycolytic products (e. g. ethanol, acetate), as for the sharper initial (up to 6 hours) biomass increase in $^{13}C$-glu to $^{12}C$-AA, most likely reflect the swap to a fresh (non-exhausted) media. Data mean±SEM of n= 4 independent cultures/per condition (culture media swap), n= 12 cultures in total. Statistics by unpaired two-sided Wilcoxon Rank Sum test; p-values are listed in **Supplementary File 3**. **c)** Fraction of intracellular $^{12}C$-AA in $^{13}C$-glu to $^{12}C$-glu and $^{13}C$-glu to $^{12}C$-AA cultures at 2h and 72h post media swap, showing the uptake of extracellular $^{12}C$-AA as indicated by the gradual increase in the intracellular fraction of $^{12}C$-AA in both cultures. Total AA levels, measured as area under the curve (AUC), were normalised per biomass, assessed by $OD_{600}$. Bar graphs represent the mean of intracellular $^{12}C$ or $^{13}C$ labelled AA fractions of n= 4 independent cultures/per condition (culture media swap), n= 8 cultures in total. Individual fraction values are listed in **Supplementary File 3**.

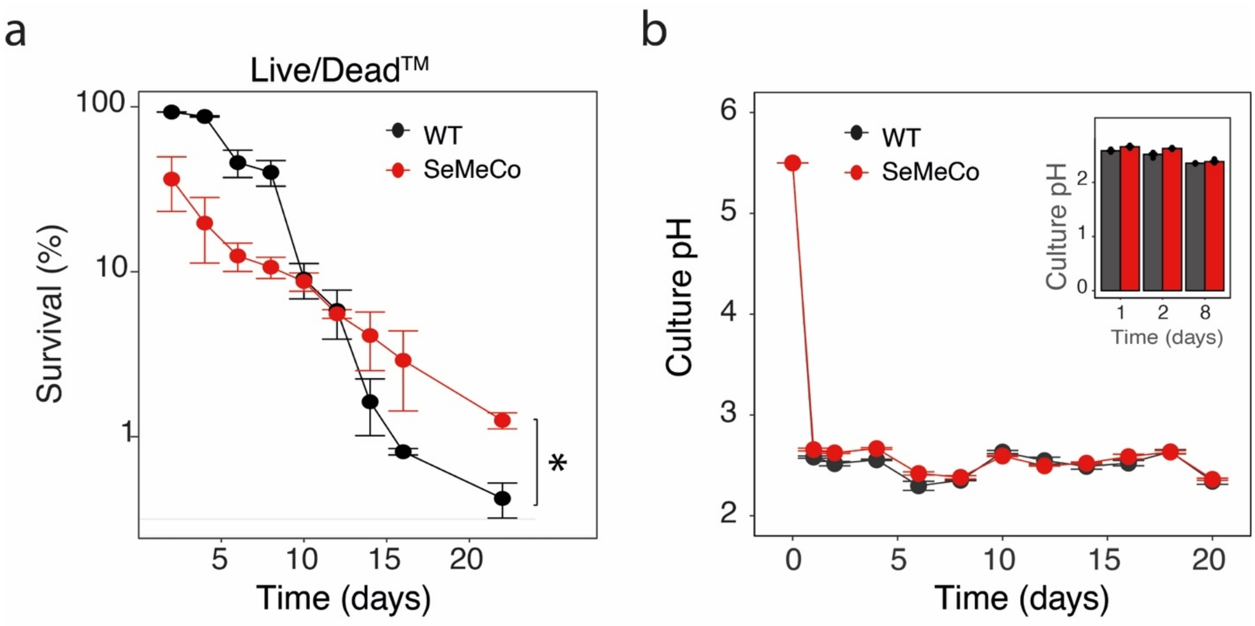


**Supplementary Fig. 4 SeMeCos survival and culture acidification during CLS. a)** Survival, as measured by staining of dead cells with the Live/Dead™ dye (see Methods) of wild-type and SeMeCos communities. Data are mean±SEM survival (percentage fold change) compared to mean wild-type survival at the beginning of stationary phase (48h culture); n= 3 independent cultures per strain. Statistics by unpaired two-sided *t*-test, p-value = 0.0108 at day 22 of culture; p-values across CLS are listed in **Supplementary File 4. b)** Culture pH values of chronologically ageing wild-type and SeMeCos communities (same cultures as in Fig. 2c). Data are mean±SEM pH values per strain; n = 4 independent cultures/ strain. Inlets are culture pH values during exponential (1 day), early stationary (2 days) and stationary (8 days) growth phases; individual dots represent independent cultures. Statistics by unpaired two-sided Wilcoxon Rank sum test; p-values are listed in **Supplementary File 5**.

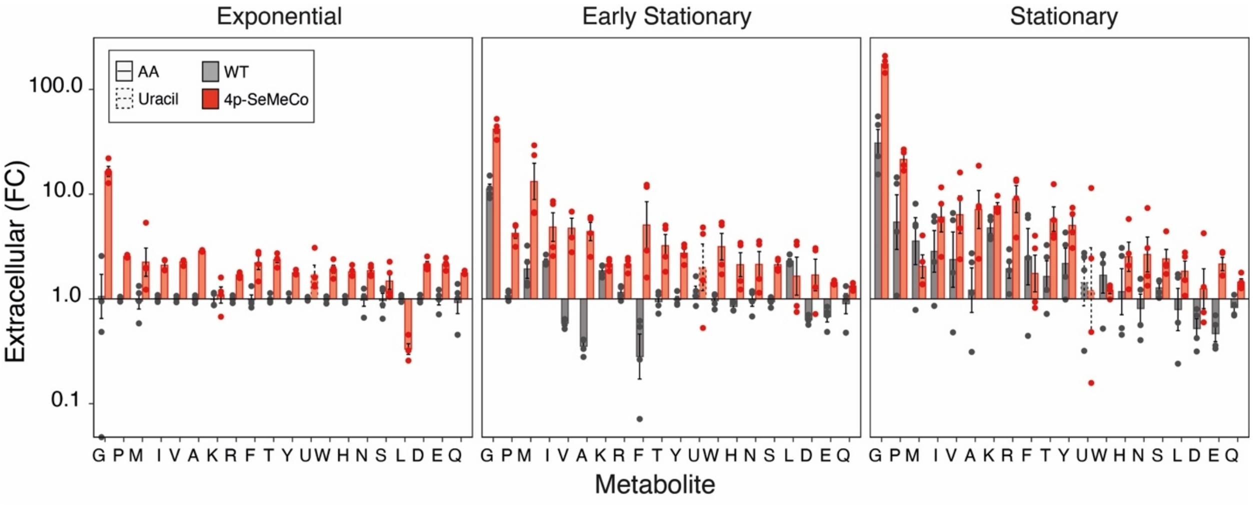


**Supplementary Fig. 5 Metabolic cooperating communities generate a rich exomebolome during chronological ageing.** Quantification of extracellular amino acids and uracil levels by targeted metabolomics (HPLC-MS/MS), in wild-type and SeMeCos communities, during exponential, early stationary and stationary growth phases. Bar plots show mean±SEM fold change (FC) to mean wild-type levels in the exponential phase of n= 4 independent cultures per strain. Concentration of each metabolite was first normalised by biomass, as assessed by $OD_{600}$. Statistics by unpaired two-sided Wilcoxon Rank Sum test and multiple testing correction using the BH method; adjusted p-values are listed in **Supplementary File 6**. Data comparing wild-type values from Fig 1c; samples were cultured, extracted and measured in parallel.

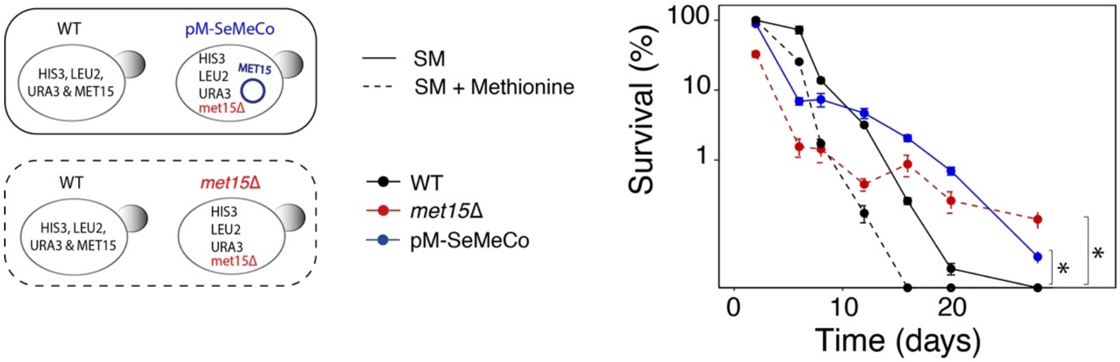


**Supplementary Fig. 6 Survival of single auxotrophs and respective single-plasmid SeMeCos for methionine.** (left) Scheme: yeast communities and respective culture conditions. 1. methionine auxotrophs (*met15Δ)* compared to wild-type yeast cultured in SM supplemented with 2g/L methionine (standard culture concentration) and 2. single-plasmid SeMeCo (pM-SeMeCo) compared to wild-type cultured in SM medium. Single auxotrophic strain (BY4741-*HIS3, LEU2, URA3*- *met15Δ*) was generated by genetic correction of 3 of the 4 auxotrophies present in the parental strain (BY4741-*his3Δ1*, *leu2Δ*, *ura3Δ, met15Δ*) with the wild-type locus *HIS3*, *LEU2* and *URA3*. The pM-SeMeCo strain was generated by complementing the *met15Δ* strain with a plasmid containing the wild-type locus *MET15*. (right) Survival of wild-type, single auxotrophs and pM-SeMeCo over time, as measured by HTP-CFU normalised to biomass. Data are mean±SEM of n= 4 independent cultures per strain per culture media condition, shown as survival (percentage fold change) compared to respective mean wild-type survival at the beginning of stationary phase (48h culture). Statistics by unpaired two-sided Wilcoxon Rank Sum test; p-value at 28 days of culture wt vs pM-SeMeCo = 3.27e-05 (SM culture) and wt vs *met15Δ* = 3.27e-05 (SM+methionine culture), p-values across CLS are listed in **Supplementary File 9**.

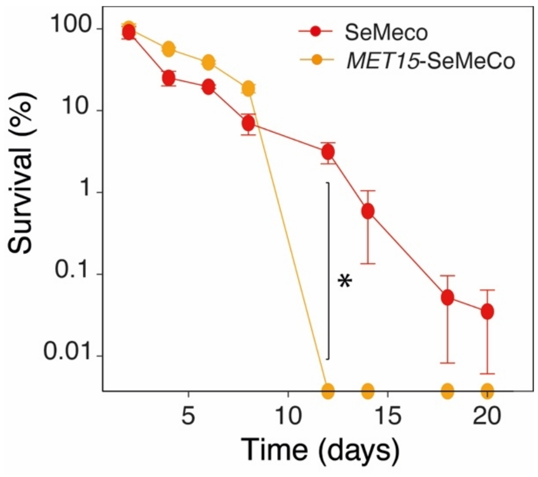


**Supplementary Fig. 7 SeMeCos where *MET15* and *met15Δ cells interact* are longer lived compared to SeMeCos unable to segregate the *MET15 locus*.** Survival percentage of SeMeCos and *MET15*-SeMeCos during CLS (CFU normalised per biomass). Data are mean±SEM survival (percentage fold change) compared to mean *MET15*-SeMeCo survival at the beginning of stationary phase (48h culture); n= 3 independent cultures per strain. Statistics by unpaired one-sided Wilcoxon Rank Sum test; p-value at day 12 of culture for SeMeCo vs *MET15*-SeMeCos = 0.031; p-values across CLS are listed in **Supplementary File 10**.

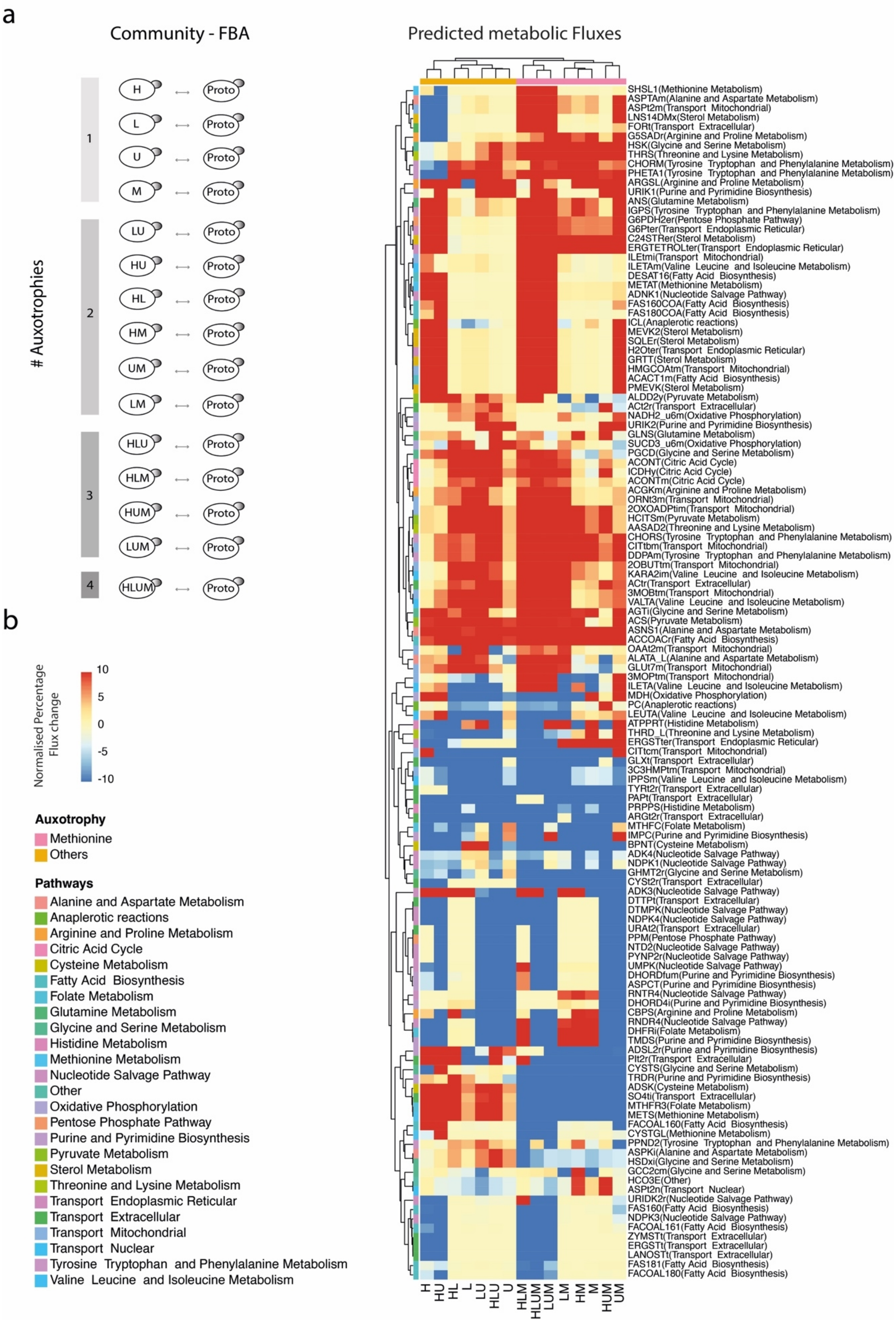
a
Community - FBA
Predicted metabolic Fluxes
### Auxotrophies
1
2
3
4
H
L
U
M
LU
HU
HL
HM
UM
LM
HLU
HLM
HUM
LUM
HLUM
Proto
SHSL1(Methionine Metabolism)
ASPTAm(Alanine and Aspartate Metabolism)
ASPt2m(Transport Mitochondrial)
LNS14DMx(Sterol Metabolism)
FORt(Transport Extracellular)
G5SADr(Arginine and Proline Metabolism)
HSK(Glycine and Serine Metabolism)
THRS(Threonine and Lysine Metabolism)
CHORM(Tyrosine Tryptophan and Phenylalanine Metabolism)
PHETA1(Tyrosine Tryptophan and Phenylalanine Metabolism)
ARGSL(Arginine and Proline Metabolism)
URIK1(Purine and Pyrimidine Biosynthesis)
ANS(Glutamine Metabolism)
IGPS(Tyrosine Tryptophan and Phenylalanine Metabolism)
G6PDH2er(Pentose Phosphate Pathway)
G6Pter(Transport Endoplasmic Reticular)
C24STRer(Sterol Metabolism)
ERGTETROLter(Transport Endoplasmic Reticular)
ILEtmi(Transport Mitochondrial)
ILETAm(Valine Leucine and Isoleucine Metabolism)
DESAT16(Fatty Acid Biosynthesis)
METAT(Methionine Metabolism)
ADNK1(Nucleotide Salvage Pathway)
FAS160COA(Fatty Acid Biosynthesis)
FAS180COA(Fatty Acid Biosynthesis)
ICL(Anaplerotic reactions)
MEVK2(Sterol Metabolism)
SQLEr(Sterol Metabolism)
H2Oter(Transport Endoplasmic Reticular)
GRTT(Sterol Metabolism)
HMGCOAtm(Transport Mitochondrial)
ACACT1m(Fatty Acid Biosynthesis)
PMEVK(Sterol Metabolism)
ALDD2y(Pyruvate Metabolism)
ACt2r(Transport Extracellular)
NADH2_u6m(Oxidative Phosphorylation)
URIK2(Purine and Pyrimidine Biosynthesis)
GLNS(Glutamine Metabolism)
SUCD3_u6m(Oxidative Phosphorylation)
PGCD(Glycine and Serine Metabolism)
ACONT(Citric Acid Cycle)
ICDHy(Citric Acid Cycle)
ACONTm(Citric Acid Cycle)
ACGKm(Arginine and Proline Metabolism)
ORNt3m(Transport Mitochondrial)
2OXOADPtim(Transport Mitochondrial)
HCITSm(Pyruvate Metabolism)
AASAD2(Threonine and Lysine Metabolism)
CHORS(Tyrosine Tryptophan and Phenylalanine Metabolism)
CITtbm(Transport Mitochondrial)
DDPAm(Tyrosine Tryptophan and Phenylalanine Metabolism)
2OBUTtm(Transport Mitochondrial)
KARA2im(Valine Leucine and Isoleucine Metabolism)
ACtr(Transport Extracellular)
3MOBtm(Transport Mitochondrial)
VALTA(Valine Leucine and Isoleucine Metabolism)
AGTi(Glycine and Serine Metabolism)
ACS(Pyruvate Metabolism)
ASNS1(Alanine and Aspartate Metabolism)
ACCOACr(Fatty Acid Biosynthesis)
OAAt2m(Transport Mitochondrial)
ALATA_L(Alanine and Aspartate Metabolism)
GLUt7m(Transport Mitochondrial)
3MOPtm(Transport Mitochondrial)
ILETA(Valine Leucine and Isoleucine Metabolism)
MDH(Oxidative Phosphorylation)
PC(Anaplerotic reactions)
LEUTA(Valine Leucine and Isoleucine Metabolism)
ATPPRT(Histidine Metabolism)
THRD_L(Threonine and Lysine Metabolism)
ERGSTter(Transport Endoplasmic Reticular)
CITtcm(Transport Mitochondrial)
GLXt(Transport Extracellular)
3C3HMPtm(Transport Mitochondrial)
IPPSm(Valine Leucine and Isoleucine Metabolism)
TYRt2r(Transport Extracellular)
PAPt(Transport Extracellular)
PRPPS(Histidine Metabolism)
ARGt2r(Transport Extracellular)
MTHFC(Folate Metabolism)
IMPC(Purine and Pyrimidine Biosynthesis)
BPNT(Cysteine Metabolism)
ADK4(Nucleotide Salvage Pathway)
NDPK1(Nucleotide Salvage Pathway)
GHMT2r(Glycine and Serine Metabolism)
CYSt2r(Transport Extracellular)
ADK3(Nucleotide Salvage Pathway)
DTTPt(Transport Extracellular)
DTMPK(Nucleotide Salvage Pathway)
NDPK4(Nucleotide Salvage Pathway)
URAt2(Transport Extracellular)
PPM(Pentose Phosphate Pathway)
NTD2(Nucleotide Salvage Pathway)
PYNP2r(Nucleotide Salvage Pathway)
UMPK(Nucleotide Salvage Pathway)
DHORDfum(Purine and Pyrimidine Biosynthesis)
ASPCT(Purine and Pyrimidine Biosynthesis)
RNTR4(Nucleotide Salvage Pathway)
DHORD4i(Purine and Pyrimidine Biosynthesis)
CBPS(Arginine and Proline Metabolism)
RNDR4(Nucleotide Salvage Pathway)
DHFRi(Folate Metabolism)
TMDS(Purine and Pyrimidine Biosynthesis)
ADSL2r(Purine and Pyrimidine Biosynthesis)
PIt2r(Transport Extracellular)
CYSTS(Glycine and Serine Metabolism)
TRDR(Purine and Pyrimidine Biosynthesis)
ADSK(Cysteine Metabolism)
SO4ti(Transport Extracellular)
MTHFR3(Folate Metabolism)
METS(Methionine Metabolism)
FACOAL160(Fatty Acid Biosynthesis)
CYSTGL(Methionine Metabolism)
PPND2(Tyrosine Tryptophan and Phenylalanine Metabolism)
ASPKi(Alanine and Aspartate Metabolism)
HSDxi(Glycine and Serine Metabolism)
GCC2cm(Glycine and Serine Metabolism)
HCO3E(Other)
ASPt2n(Transport Nuclear)
URIDK2r(Nucleotide Salvage Pathway)
FAS160(Fatty Acid Biosynthesis)
NDPK3(Nucleotide Salvage Pathway)
FACOAL161(Fatty Acid Biosynthesis)
ZYMSTt(Transport Extracellular)
ERGSTt(Transport Extracellular)
LANOSTt(Transport Extracellular)
FAS181(Fatty Acid Biosynthesis)
FACOAL180(Fatty Acid Biosynthesis)
H
HU
HL
L
LU
HLU
U
HLM
HLUM
LUM
LM
HM
M
HUM
UM
b
Normalised Percentage Flux change
10
5
0
-5
-10
Auxotrophy
Methionine
Others
Pathways
Alanine and Aspartate Metabolism
Anaplerotic reactions
Arginine and Proline Metabolism
Citric Acid Cycle
Cysteine Metabolism
Fatty Acid Biosynthesis
Folate Metabolism
Glutamine Metabolism
Glycine and Serine Metabolism
Histidine Metabolism
Methionine Metabolism
Nucleotide Salvage Pathway
Other
Oxidative Phosphorylation
Pentose Phosphate Pathway
Purine and Pyrimidine Biosynthesis
Pyruvate Metabolism
Sterol Metabolism
Threonine and Lysine Metabolism
Transport Endoplasmic Reticular
Transport Extracellular
Transport Mitochondrial
Transport Nuclear
Tyrosine Tryptophan and Phenylalanine Metabolism
Valine Leucine and Isoleucine Metabolism

**Supplementary Fig. 8 Genome-scale metabolic modelling clustering of metabolic pathway fluxes according to the presence or absence of *met15Δ cells in a community*. a)** Genome-scale metabolic modelling, using flux balance analysis (FBA), on pairwise communities of a prototroph (wild-type) and each of the possible auxotrophs in a SeMeCo, in a total of 15 communities pairs analysed from [1]. **b)** Heatmap showing predicted metabolic pathway flux change, clustering according to interactions with methionine auxotrophs (unsupervised clustering). Columns are each prototroph-auxotroph community (auxotrophic label shown) and rows are metabolic pathways. Predicted metabolic fluxes are represented in a -10 to 10 scale, where absolute percentage flux changes in auxotrophs vs prototroph >10% are scaled to 10 or -10 according to the directionality of change, as increased or decreased respectively (**Supplementary File 12**).

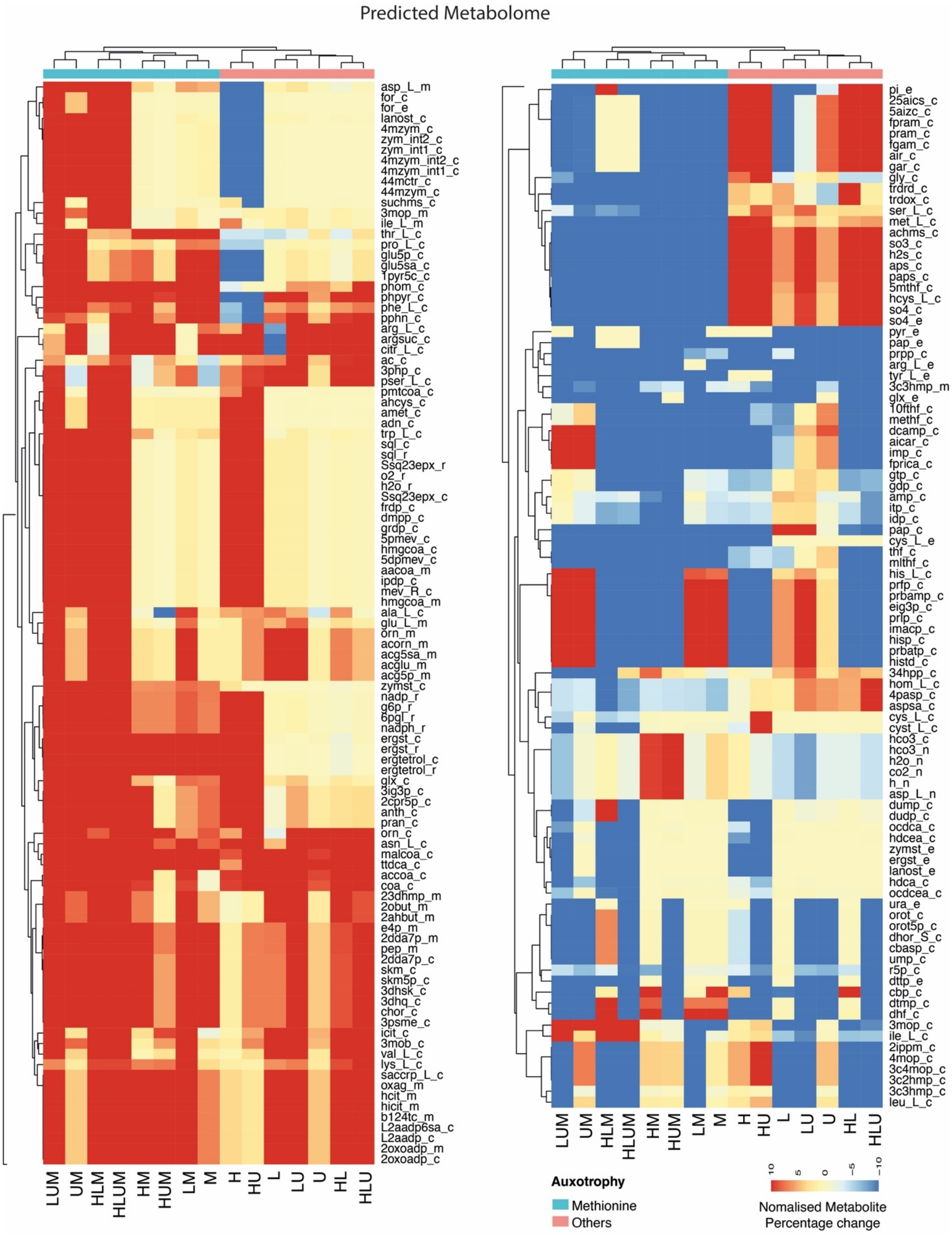
Predicted Metabolome
asp_L_m
for_c
for_e
lanost_c
4mzym_c
zym_int2_c
zym_int1_c
4mzym_int2_c
4mzym_int1_c
44mctr_c
44mzym_c
suchms_c
3mop_m
ile_L_m
thr_L_c
pro_L_c
glu5p_c
glu5sa_c
1pyr5c_c
phom_c
phpyr_c
phe_L_c
pphn_c
arg_L_c
argsuc_c
citr_L_c
ac_c
3php_c
pser_L_c
pmtcoa_c
ahcys_c
amet_c
adn_c
trp_L_c
sql_c
sql_r
Ssq23epx_r
o2_r
h2o_r
Ssq23epx_c
frdp_c
dmpp_c
grdp_c
5pmev_c
hmgcoa_c
5dpmev_c
aacoa_m
ipdp_c
mev_R_c
hmgcoa_m
ala_L_c
glu_L_m
orn_m
acorn_m
acg5sa_m
acglu_m
acg5p_m
zymst_c
nadp_r
g6p_r
6pgl_r
nadph_r
ergst_c
ergst_r
ergtetrol_c
ergtetrol_r
glx_c
3ig3p_c
2cpr5p_c
anth_c
pran_c
orn_c
asn_L_c
malcoa_c
ttdca_c
accoa_c
coa_c
23dhmp_m
2obut_m
2ahbut_m
e4p_m
2dda7p_m
pep_m
2dda7p_c
skm_c
skm5p_c
3dhsk_c
3dhq_c
chor_c
3psme_c
icit_c
3mob_c
val_L_c
lys_L_c
saccrp_L_c
oxag_m
hcit_m
hicit_m
b124tc_m
L2aadp6sa_c
L2aadp_c
2oxoadp_m
2oxoadp_c
LUM
UM
HLM
HLUM
HM
HUM
LM
M
H
HU
L
LU
U
HL
HLU
pi_e
25aics_c
5aizc_c
fpram_c
pram_c
fgam_c
air_c
gar_c
gly_c
trdrd_c
trdox_c
ser_L_c
met_L_c
achms_c
so3_c
h2s_c
aps_c
paps_c
5mthf_c
hcys_L_c
so4_c
so4_e
pyr_e
pap_e
prpp_c
arg_L_e
tyr_L_e
3c3hmp_m
glx_e
10fthf_c
methf_c
dcamp_c
aicar_c
imp_c
fprica_c
gtp_c
gdp_c
amp_c
itp_c
idp_c
pap_c
cys_L_e
thf_c
mlthf_c
his_L_c
prfp_c
prbamp_c
eig3p_c
prlp_c
imacp_c
hisp_c
prbatp_c
histd_c
34hpp_c
hom_L_c
4pasp_c
aspsa_c
cys_L_c
cyst_L_c
hco3_c
hco3_n
h2o_n
co2_n
h_n
asp_L_n
dump_c
dudp_c
ocdca_c
hdcea_c
zymst_e
ergst_e
lanost_e
hdca_c
ocdcea_c
ura_e
orot_c
orot5p_c
dhor_S_c
cbasp_c
ump_c
r5p_c
dttp_e
cbp_c
dtmp_c
dhf_c
3mop_c
ile_L_c
2ippm_c
4mop_c
3c4mop_c
3c2hmp_c
3c3hmp_c
leu_L_c
LUM
UM
HLM
HLUM
HM
HUM
LM
M
H
HU
L
LU
U
HL
HLU
Auxotrophy
Methionine
Others
10
5
0
−5
−10
Nomalised Metabolite
Percentage change

**Supplementary Fig. 9 Genome-scale metabolic modelling predicts specific metabolome changes in communities where *MET15* and *met15Δ cells interact*.** Heatmap showing predicted metabolite clustering according to when prototrophs in a community interact with methionine auxotrophs (unsupervised clustering). Columns are each prototroph-auxotroph community (auxotrophic label shown) and rows are predicted metabolites. Predicted metabolites are represented in a -10 to 10 scale, where absolute percentage metabolite changes in auxotrophs vs prototroph >10% are scaled to 10 or -10 according to the directionality of change, as increased or decreased respectively. The heatmap was split for visualisation. Full list of metabolite names is in **Supplementary File 12**.

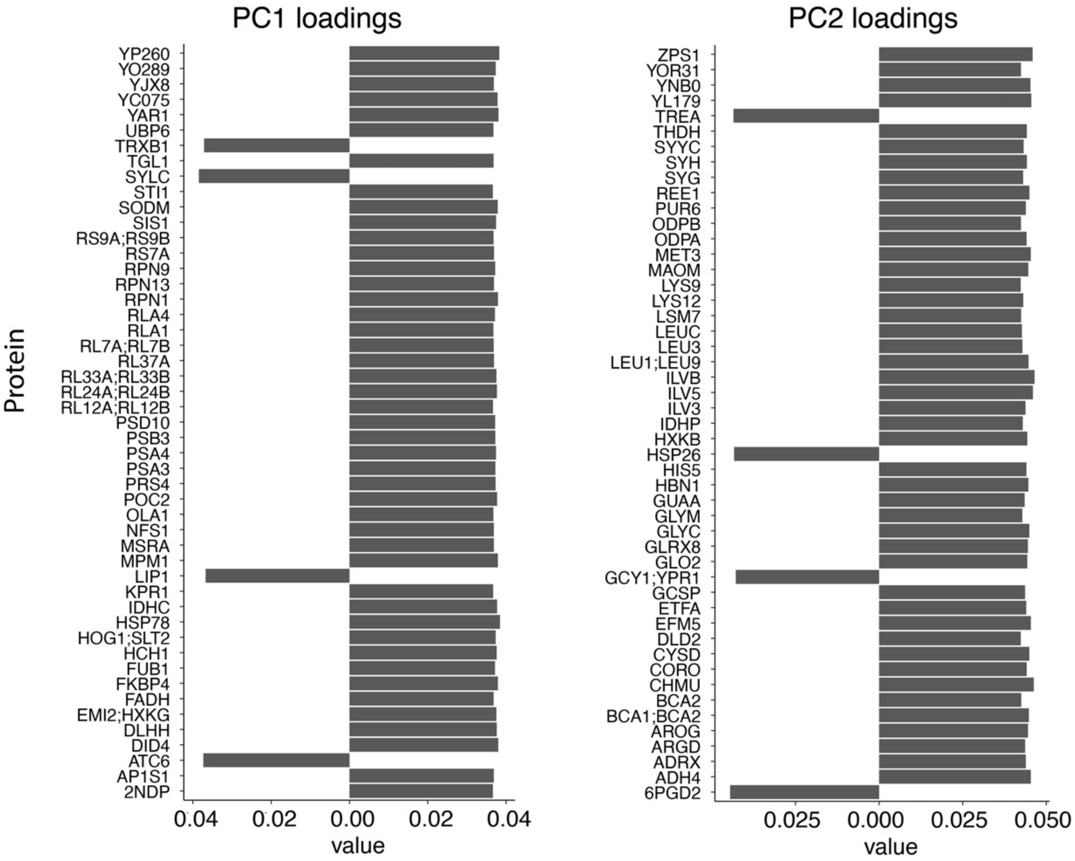


**Supplementary Fig. 10 Principal component analysis demonstrates that major proteome changes are associated with transition from exponential to stationary phase and methionine cooperation.** Top 50 loadings from PCA1 and PCA2 mostly comprising proteins associated with growth phase and metabolic changes respectively. Analysis derived from PCA analysis in Fig. 4b.

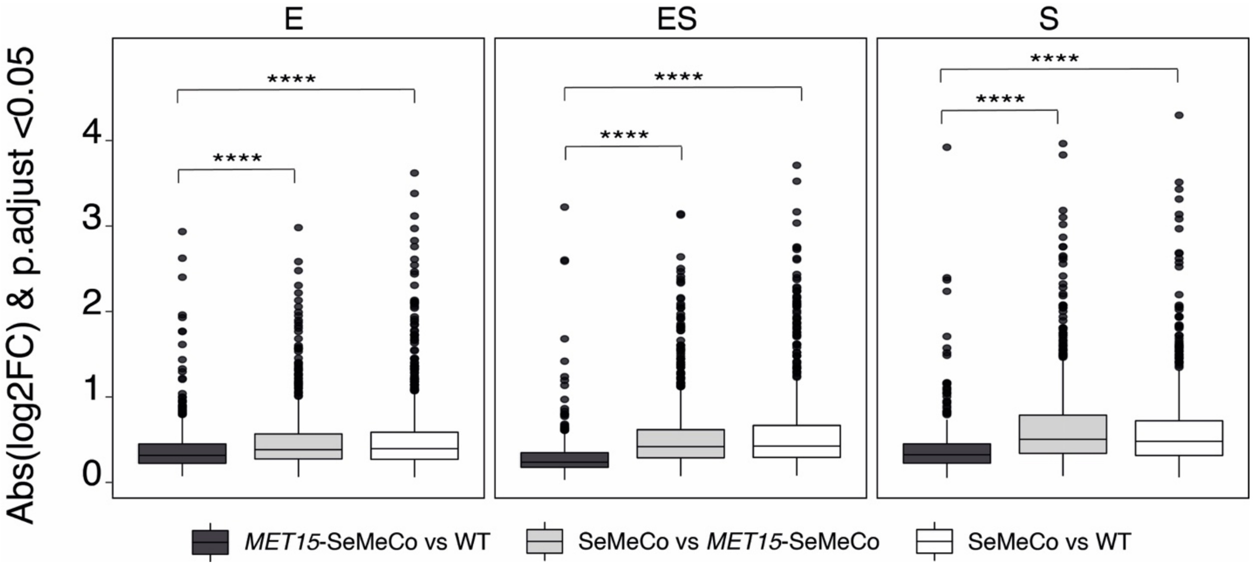


**Supplementary Fig. 11 Higher magnitude of proteome changes in communities where *MET15* and *met15Δ cells interact*.** SeMeCos, *MET15*-SeMeCos and control wild-type cultures were collected at exponential (E), early stationary (ES) and stationary (S) growth phases, for proteomics analysis using micro-flow LC-SWATH MS [2] and DIA-NN [3] as described in Fig. 4a. Proteomics performed in 4 independent cultures (biological replicates) per yeast strain (total n=12 cultures). Box plots showing absolute log2 fold change (Abs(log2FC)) of significant differentially expressed proteins (DEP, adjusted p-value <0.05), in pairwise comparisons of the different yeast communities according to the proteomics data. Box plots represent median (50% quantile, middle line) and lower and upper quantiles (lower (25% quantile) and upper (75% quantile). Statistics by unpaired two-sided *t*-test and multiple testing correction using the BH method; adjusted p-value **** < 0.00005.

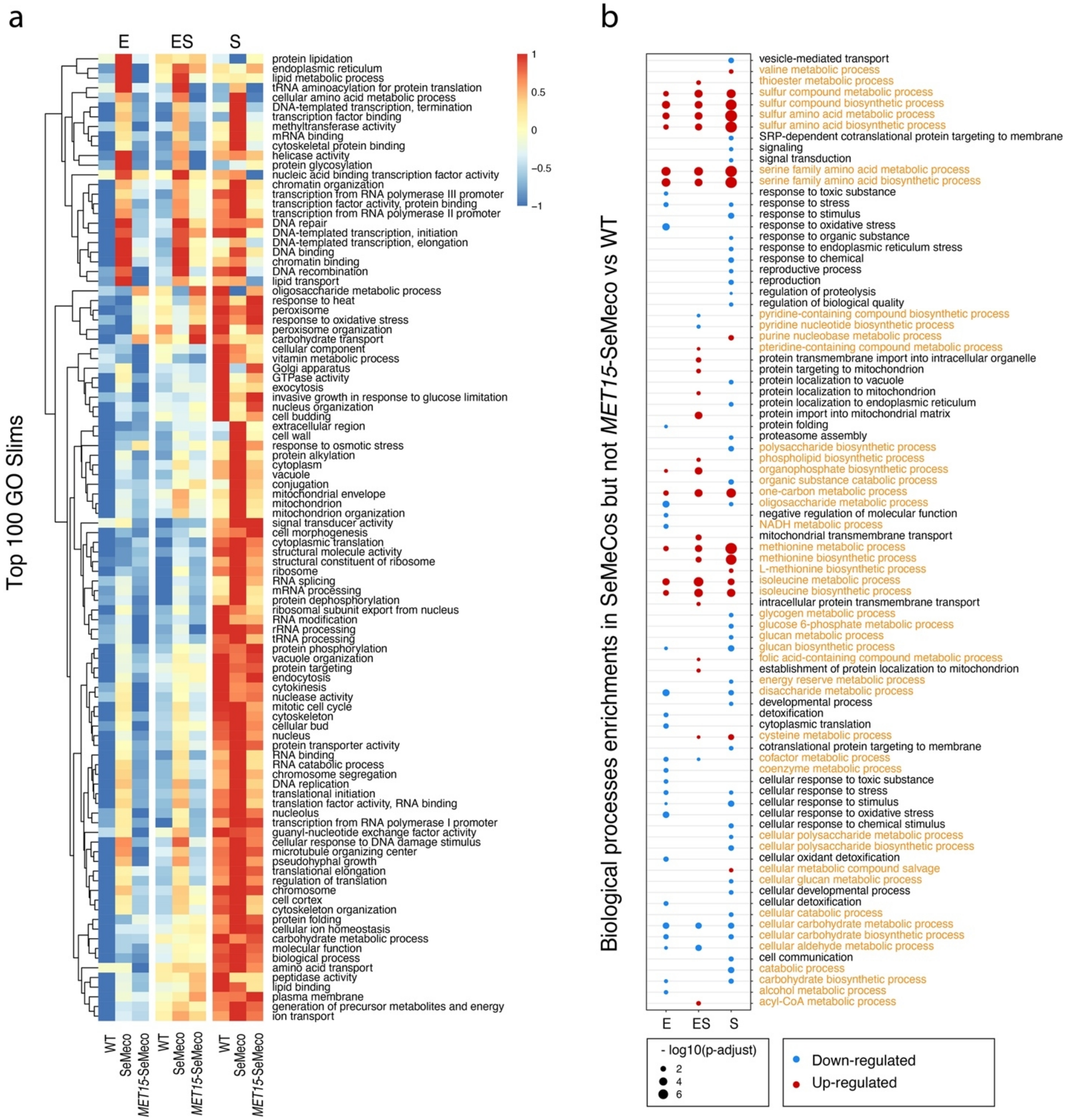


**Supplementary Fig. 12 Widespread proteome and metabolome changes in yeast communities where *MET15* and *met15Δ* cells interact. a)** Heatmap derived from proteome analysis in Fig. 4a, showing expression of proteins, normalised to a -1 to 1 scale, belonging to top 100 GO slim terms (raws), per growth phase and yeast communities (columns). **b)** Gene set enrichment analysis (GSEA) showing significantly enriched (BH adjusted p-value cutoff <0.05) gene ontologies (GO) comprising biological processes, as defined in the KEGG database, in SeMeCos, but not in *MET15*-SeMeCos, compared to wild-type (WT) communities, during exponential (E), early stationary (ES) and stationary (S) growth phases. Data are from the proteomics analysis as described in Fig. 4a. Statistics by unpaired two-sided *t*-test and multiple testing correction using the BH method; adjusted-p-values are listed in **Supplementary File 14**.

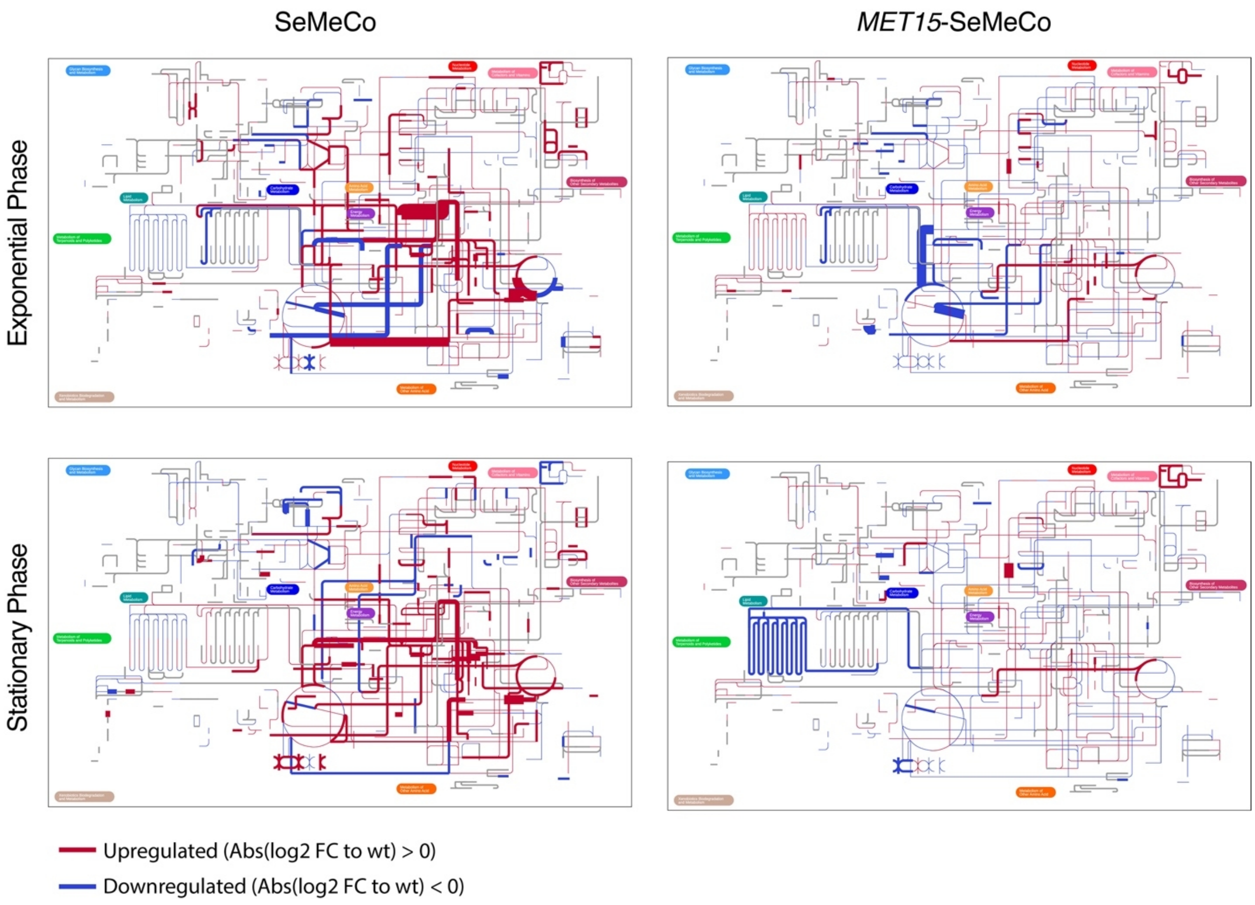


**Supplementary Fig. 13 Metabolic interactions between *MET15* and *met15Δ* cells induce widespread metabolic changes.** Differential metabolic enzyme expression levels, derived from the proteome analysis in Fig. 4a, mapped to the yeast metabolic network using IPATH3 [4] in exponential and stationary phases. Red and blue lines represent significantly (BH adjusted p-value <0.05) up- or down-regulated proteins in SeMeCos and *MET15*-SeMeCos when compared to wild-type; grey lines represent non-mapped/absent proteins in the measured proteomes. Thickness of the lines represent absolute log2 fold change (Abs(log2FC)) changes (thickening = increased Abs(log2FC)).

**Supplementary Table S1: yeast strains**

| Name | Species/strain | Source | Genotype |
|---|---|---|---|
| Parental | *S.cerevisiae, BY4741* | ATCC | *His3Δ, leu2Δ, ura3Δ, met15Δ* |
| Prototrophic | *S.cerevisiae, BY4741* | Yu & Correia-Melo *et al.* 2022 [1] | *HIS3, LEU2, URA3, MET15* |
| SeMeCo | *S.cerevisiae, BY4741* | Yu & Correia-Melo *et al.* 2022 [1] | *pHIS3, pLEU2, pURA3, pMET15* |
| *HIS3*-SeMeCo | *S.cerevisiae, BY4741* | This study | *HIS3, pLEU2, pURA3 , pMET15* |
| *LEU2*-SeMeCo | *S.cerevisiae, BY4741* | This study | *LEU2, pHIS3, pURA3 , pMET15* |
| *URA3*-SeMeCo | *S.cerevisiae, BY4741* | This study | *URA3, pHIS3, pLEU2, pMET15* |
| *MET15*-SeMeCo | *S.cerevisiae, BY4741* | This study | *MET15, pHIS3, pLEU2, pURA3* |
| *met15Δ* | *S.cerevisiae, BY4741* | This study | *HIS3, LEU2, URA3, met15Δ* |
| pM-SeMeCo | *S.cerevisiae, BY4741* | This study | *HIS3, LEU2, URA3, pMET15* |

**Supplementary Table S2: primers for genomic integration of the metabolic markers**

| Metabolic marker | Primer Fw | Primer Rv |
|---|---|---|
| *HI3* | 5' TATCGTTTGAACACGGCATT 3' | 5' CGCGCCTCGTTCAGAATGAC 3' |
| *LEU2* | 5' GAATTAAAGGATTGGATAGC 3' | 5' CCCTATGAACATATTCCATT 3' |
| *URA3* | 5' GTTCATCATCTCATGGATCT 3' | 5' TACTGTTACTTGGTTCTGGC 3' |
| *MET15* | 5' TCGTTTTCTACTTTCTTCTG 3' | 5' GGAGAAGTCAAGACTATGAA 3' |

*Note: Primers were designed to flank the genomic region of the gene of interest ± several hundred base pairs to maximise the homologous region for recombination. The YSBN5 yeast strain was used as template DNA.*

#### Supplementary Table S3: plasmid for SeMeCo generation

| Metabolic marker | Addgene | Reference |
|---|---|---|
| *pHIS3* | #64178 | Mülleder *et al.* 2016 [5] |
| *pLEU2* | #64177 | Mülleder *et al.* 2016 [5] |
| *pURA3* | #64180 | Mülleder *et al.* 2016 [5] |
| *pMET15* | #64179 | Mülleder *et al.* 2016 [5] |
